# Deletion of the microglial transmembrane immune signaling adaptor TYROBP ameliorates Huntington’s disease mouse phenotype

**DOI:** 10.1101/2022.02.18.480944

**Authors:** Jordi Creus-Muncunill, Daniele Mattei, Joanna Bons, Angie V. Ramirez-Jimenez, B. Wade Hamilton, Chuhyon Corwin, Sarah Chowdhury, Birgit Schilling, Lisa Ellerby, Michelle E. Ehrlich

**Affiliations:** Department of Neurology, Icahn School of Medicine at Mount Sinai, New York, USA; Nash Family Department of Neuroscience and Friedman Brain Institute, Icahn School of Medicine at Mount Sinai, New York, USA; Buck Institute for Research on Aging, Novato, California, USA

## Abstract

**BACKGROUND:** Huntington’s disease (HD) is a fatal neurodegenerative disorder caused by an expansion of the CAG trinucleotide repeat in the huntingtin gene. Immune activation is abundant in the striatum of HD patients. Detection of active microglia at presymptomatic stages suggests that microgliosis is a key early driver of neuronal dysfunction and degeneration. Recent studies showed that deletion of *Tyrobp*, a microglial-enriched protein, ameliorates neuronal function in Alzheimer’s disease amyloid and tauopathy mouse models while decreasing components of the complement subnetwork, thus raising the possibility that *Tyrobp* deletion can be beneficial for HD.

**METHODS:** To test the hypothesis that *Tyrobp* deficiency would be beneficial in a HD model, we placed the Q175 HD mouse model on a *Tyrobp*-null background. We characterized these mice with a combination of behavioral testing, immunohistochemistry, transcriptomic and proteomic profiling. Further, we evaluated the Q175 microglia-specific gene signature, with and without *Tyrobp*, by purifying microglia from these mice for transcriptomics.

**RESULTS:** Comprehensive analysis of publicly available transcriptomic HD human data revealed that TYROBP network is overactivated in HD putamen. The Q175 mice showed morphologic microglial activation, reduced levels of post-synaptic density-95 protein and motor deficits at 6 and 9 months of age, all of which were ameliorated on the *Tyrobp*-null background. Gene expression analysis revealed that lack of *Tyrobp* in the Q175 model does not prevent the decrease in the expression of striatal neuronal genes but reduces pro-inflammatory pathways that are specifically active in HD human brain. Integration of transcriptomic and proteomic data identified that astrogliosis and complement system pathway were reduced after *Tyrobp* deletion. Results were further validated by immunofluorescence analysis. Microglia-specific HD gene dysregulation, characterized by overexpression of neuronal genes, was also not corrected by *Tyrobp* deletion.

**CONCLUSIONS:** Our data provide molecular and functional support demonstrating that *Tyrobp* deletion prevents many of the abnormalities in the Q175 HD mouse model, in the absence of correction of striatal neuronal gene expression, suggesting that the *Tyrobp* pathway is a potential therapeutic candidate for Huntington’s disease.

## BACKGROUND

Huntington’s disease (HD) is a neurodegenerative disease caused by an expansion of the trinucleotide CAG within exon-1 of the Huntingtin (*HTT*) gene. The resultant protein (mutant Huntingtin; mHtt) contains an aberrant polyglutamine tail to which neurotoxicity is attributed [1]. HD is characterized by a progressive loss of striatal projection neurons, known as medium spiny neurons (MSNs) [2,3], which leads to motor alterations, cognitive deficits, and eventual death. Neurodegeneration, however, is not solely restricted to the striatum since cortical atrophy is also evident in HD patients [4,5]. There are several cell autonomous and non-cell autonomous mechanisms by which the *HTT* mutation causes disease [6], including a role for glial cells in both the onset and progression of the disease [7]. Huntingtin (Htt) is highly expressed in microglia and mHtt aggregates are accumulated in this cell type [8]. Brains from presymptomatic HD human carriers contain activated microglia, which have been identified as morphologically abnormal, with increased size, amoeboid-like cell body shape, short or absent processes and, increased phagocytic activity [9–11]. Accordingly, there is an increase of pro-inflammatory cytokines, including IL-1β, in HD individuals [12–14]. Transcriptional profiling of HD brains revealed that pro-inflammatory pathways are activated in the most affected brain regions, i.e. caudate/putamen and cortex [15–17]. HD mouse models largely fail to recapitulate the activation of pro-inflammatory pathways at a transcriptional level [18]. However, restricted expression of mHtt in microglia is sufficient to induce both pro-inflammatory gene expression and inflammation, with an associated exacerbation of neurodegeneration [19]. These studies have firmly established the contribution of mHtt within microglial cells as a driver of HD, and have spurred the development of immunomodulatory therapeutic approaches, which are currently being tested in the clinic.

We previously determined that deletion of *Tyrobp* (also known as DAP12) in Alzheimer’s disease (AD) amyloid and tauopathy mouse models mitigates cognitive dysfunction and restores neuronal function [20–22]. TYROBP is a microglial-enriched, phosphotyrosine phosphoprotein that has been genetically associated with late onset Alzheimer’s disease (LOAD) [23]. Mechanistically, TYROBP works in protein complexes acting as the adaptor for proteins such as TREM2 and the complement receptor CR3 to regulate phagocytosis and signal transduction [24–26]. Genetic ablation of *Tyrobp* in amyloidosis and tauopathy models decreases microglial clustering around amyloid plaques, reduces members of the classical complement pathway, and represses the induction of inflammatory-related genes related to disease-associated microglia [20,21]. In this study, we investigate the deletion of *Tyrobp* as a potential therapeutic strategy in HD, asking whether knocking out *Tyrobp*, a contributor to reactive microglia, is protective in a mouse model of the disease.

## RESULTS

### *TYROBP* network is overactivated in human HD putamen

Down-regulated genes in HD are known to be largely neuronal, and in the striatum, expressed in medium spiny neurons. To determine which pathways are activated in HD brains, we performed a comprehensive review of publicly available human gene expression datasets, with a focus on upregulated differentially expressed genes (DEGs). We observed a distinct pattern of gene expression that is consistent across multiple datasets, with upregulated genes dominated by those involved in immune and inflammatory-related processes. RNAseq-derived datasets show clear enrichment for inflammatory pathways and importantly, *TYROBP* network in microglia. Microarray datasets revealed DEGs associated with Hedgehog signaling pathway, EGF/EGFR Signaling Pathway and focal adhesion, which include integrin-encoding genes, possibly related to microglial motility and activation (Fig. 1A). Enrichment analysis results obtained from each dataset are shown in Supplementary Table 1. We next identified the genes that are commonly upregulated across all HD RNAseq datasets, resulting in a total of 221 genes. To inquire into the role of commonly upregulated DEGs across HD brains, we analyzed them by means of the STRING v11.5 tool. The analysis generated a gene/protein interaction network (Fig. 1B; strength of interaction score > 0.7) consisting of 217 DEGs (nodes) and 99 edges. Nodes devoid of any interactions were deleted from the network. With 8 edges, *TYROBP* is the most interactive node, followed by *RELA, SYK* and *TGFB1* with 7 edges each. GO terms associated with the network are related to response to stimulus and immune system process (Supplementary Table 2). Functionally related significant modules from the network were mined with MCODE with cutoffs of the MCODE score ≥2 and number of nodes ≥2. Three significant modules were detected, and the top module with MCORE score 4.5 (5 nodes, 9 edges) was formed by metallothionein-related proteins. The second most significant module with MCORE score 4.0 (4 nodes, 6 edges) contained *TYROBP, SYK, FGR* and *FCGR2A*. Interestingly, TYROBP signals through SYK to activate intracellular pathways including extracellular signal-regulated protein kinase (ERK), phosphatidylinositol 3-kinase (PI3K), phospholipase Cγ (PLCγ), and Vav [27]. Together, these data suggest that *TYROBP* may play an important role in the pathophysiologic process in HD.

**Fig. 1.**
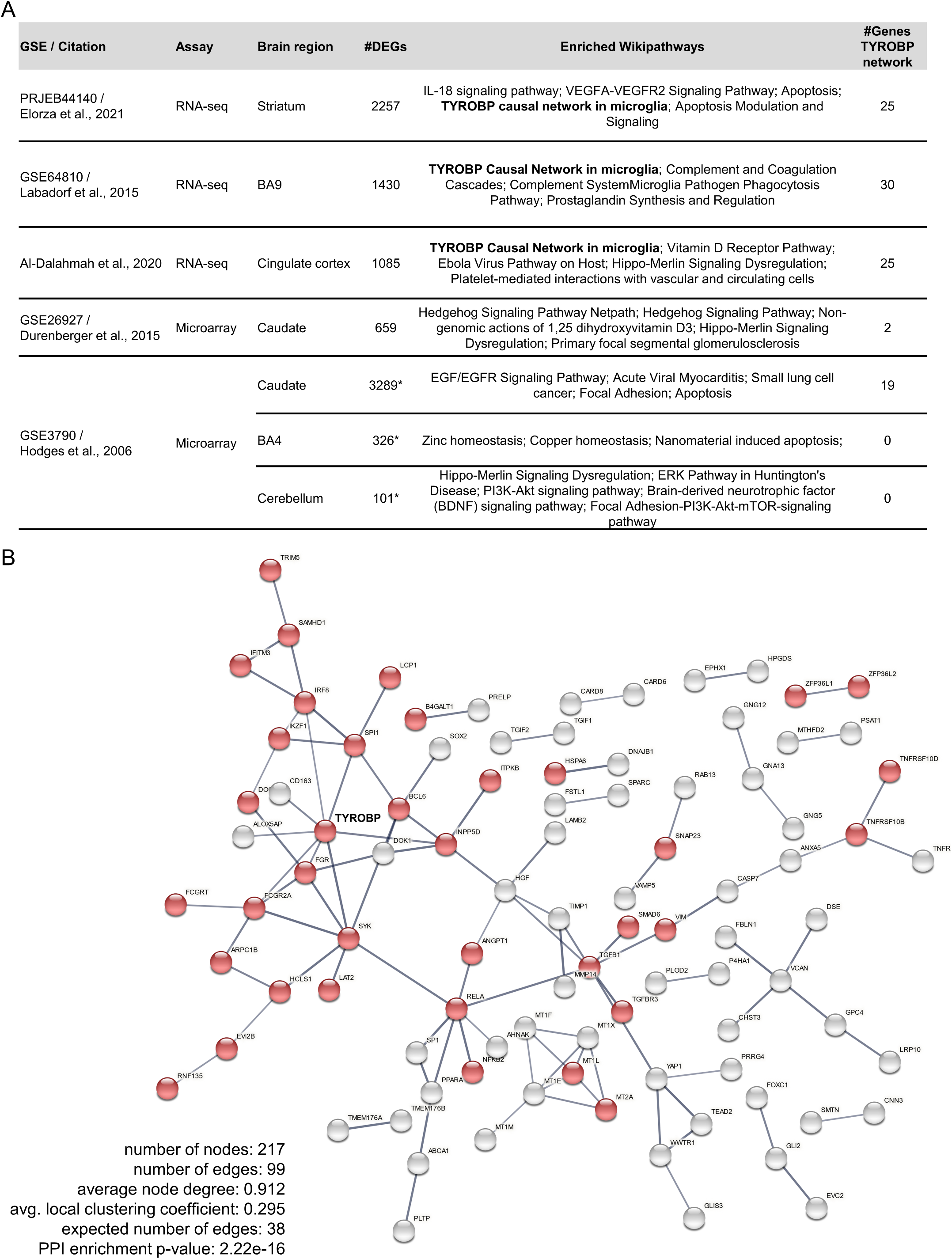
TYROBP network is upregulated in HD brains. A. Overview of the published datasets used in this study. *nominal p-value < 0.001 was used as threshold. B. STRING-generated interaction network of the common differentially expressed genes (DEGs) detected by RNA-seq in HD brains. STRING v11.5 was used to derive the network of 221 DEGs applying following prediction methods: text mining, experiments, databases, co-expression, neighborhood, gene fusion, co-occurrence. The nodes that did not interact with other nodes were deleted. The red nodes illustrate gene/proteins assigned to “immune system process” Gene Ontology pathway. Line thickness indicates the strength of data support. Interaction score > 0.7.

### *Tyrobp* deletion prevents pro-inflammatory phenotype of Q175 microglia

We previously reported that deletion of *Tyrobp* altered microglial response to AD-related pathologies, including amyloidosis and tauopathy [20–22]. Microgliosis and neuroinflammation are evident in HD human caudate/putamen beginning at presymptomatic stages [10]. Our in-silico analysis pointed to a key role of *TYROBP* in HD pathophysiology, so we performed a mouse genetics analysis, taking advantage of a previously generated *Tyrobp*-deficient [*Tyrobp* homozygous knockout (*Tyrobp*^*(-/-)*^)] mouse line [28]. *Tyrobp*^*(-/-)*^ mice were bred with Q175 heterozygous mice to analyze Q175 mice on a *Tyrobp*^*(-/-)*^ background. Q175 model is a genetically precise mouse model of adult-onset HD, in which human *HTT* exon (with a ∼190 CAG repeats) was knocked into the endogenous *Htt*. This well-defined model recapitulates many of the molecular, neuropathological, and behavioral abnormalities observed in HD patients. We evaluated the effects of *Tyrobp* deletion on microglial number in Q175 mice at 10 months of age, a fully symptomatic time-point. We detected an increase in the number of microglial cells in the striatum of Q175 mice, independent of *Tyrobp* expression (Fig. 2A). We did not detect changes in Iba1 fluorescence intensity (Fig. 2B), nor mRNA or protein levels (Supplementary Fig. 1). Next, we analyzed microglial morphology.

**Fig. 2.**
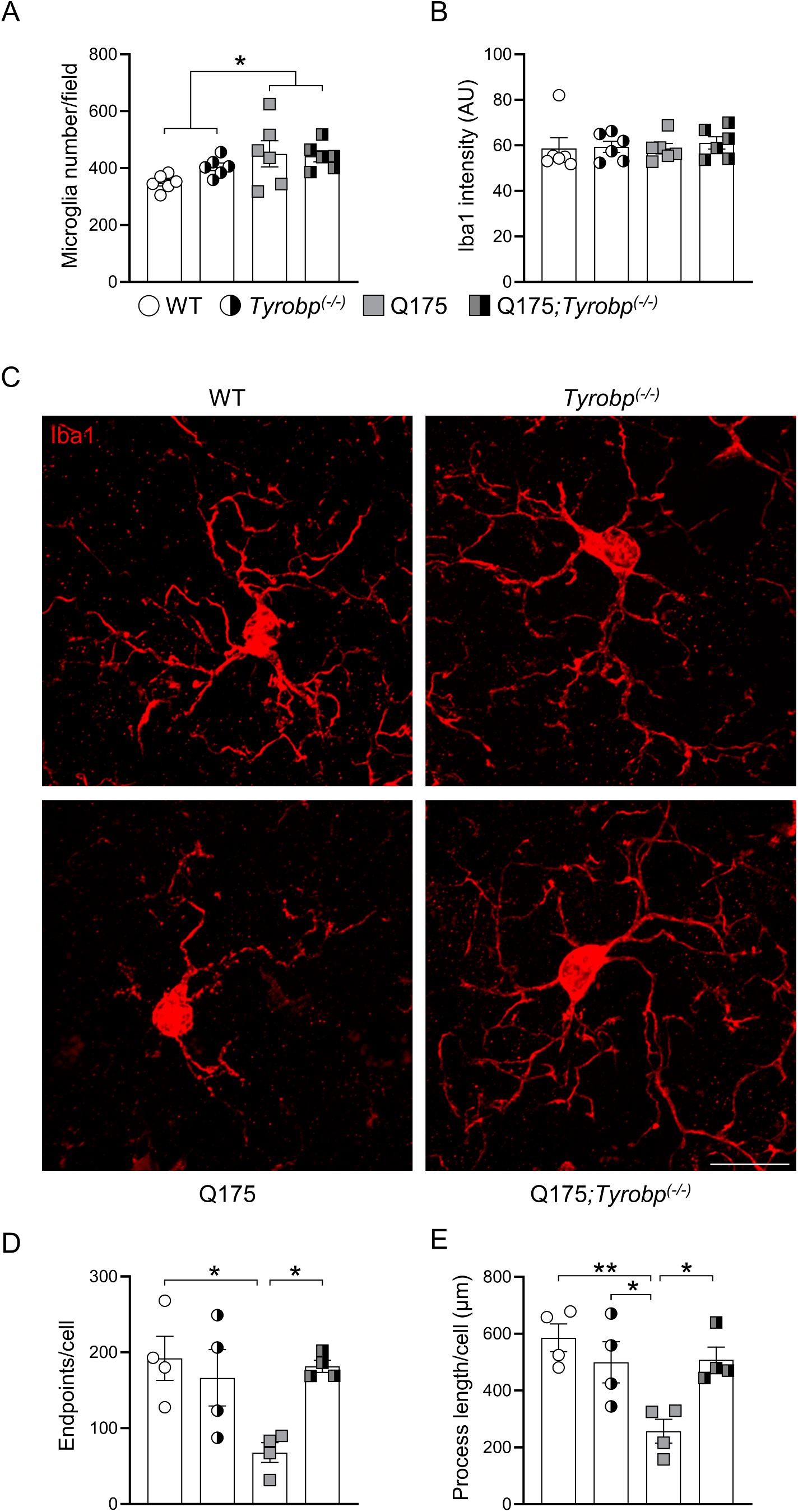
*Tyrobp* deletion corrects microglia morphologic parameters. Quantification of Iba1-immunopositive (A) cell number and (B) intensity in striatal area (n = 6 mice per group with an average of 3 slices per mouse). (C) Illustrative Iba-1 immunostaining images of the striatum from 10-month old WT and Q175 mice with (left) and without (right) *Tyrobp*. Scale bar: 15 µm. (D) Endpoints/cell and (E) process length of striatal microglia. n = 15 microglia per mouse with n = 4 mice per group. Error bars represent means ± SEM. Males and females were used for experiments, and results were combined for analysis. Statistical analyses were performed using a Two-Way ANOVA with Bonferroni as post-hoc test. *p < 0.05; **p < 0.01.

Microglial cells from Q175 mice showed decreased branch numbers and total branch length (Fig. 2C-E), indicative of a pro-inflammatory status [29]. Importantly, microglia morphology from Q175;*Tyrobp*^*(-/-)*^ mice was similar to WT microglia (Fig. 2C-E). These data show that *Tyrobp* deletion in Q175 mice prevents HD-associated microglia morphological changes.

### *Tyrobp* deletion reduces microglia CD68 content and impedes PSD-95 loss in Q175 mice

Active microglia are characterized by increased phagocytic activity [30]. CD68 is a transmembrane protein highly expressed in phagocytic lysosomes of microglial cells [31]. High levels of CD68 have been detected in the R6/2 mice, a transgenic mouse model of HD with rapid progression, beginning at early disease stages [32]. To determine the functional implications of the improved morphological indices in Q175;*Tyrobp*^*(-/-)*^ mice, we evaluated the area covered by CD68 in striatal microglia. *Tyrobp* deletion in Q175 mice normalized striatal CD68 content (Fig. 3A, B). Striatal CD68 mRNA levels were not altered, suggesting that CD68 levels are modulated through post-transcriptional mechanisms (Fig. 3C). Increased CD68 content in Q175 may point to an aberrant excess of phagocytosis or, in contrast, a reflection of an overwhelmed phagolysosome system incapable of processing the phagocytosed material. In many neurodegenerative diseases, including HD, aberrant microglia engulf and eliminate synapses in a non-physiological manner, leading to synaptic deficits [33,34]. Here, we evaluated post-synaptic density 95 (PSD-95) protein levels and estimated the density of PSD-95 immunopositive puncta in striatum in our four groups of mice. In agreement with previous reports [35–38], we detected reduced PSD-95 protein levels in the striatum of Q175 mice (Fig. 3D). Importantly, deletion of *Tyrobp* in Q175 mice prevented PSD-95 protein reduction (Fig. 3D). Accordingly, immunostaining revealed that *Tyrobp* deletion in Q175 mice prevented PSD-95 puncta reduction (Fig. 3E). These data support an interpretation that microglial phagocytic activity is reduced in HD mice in the absence of *Tyrobp*, leading to a preservation of synapses.

**Fig. 3.**
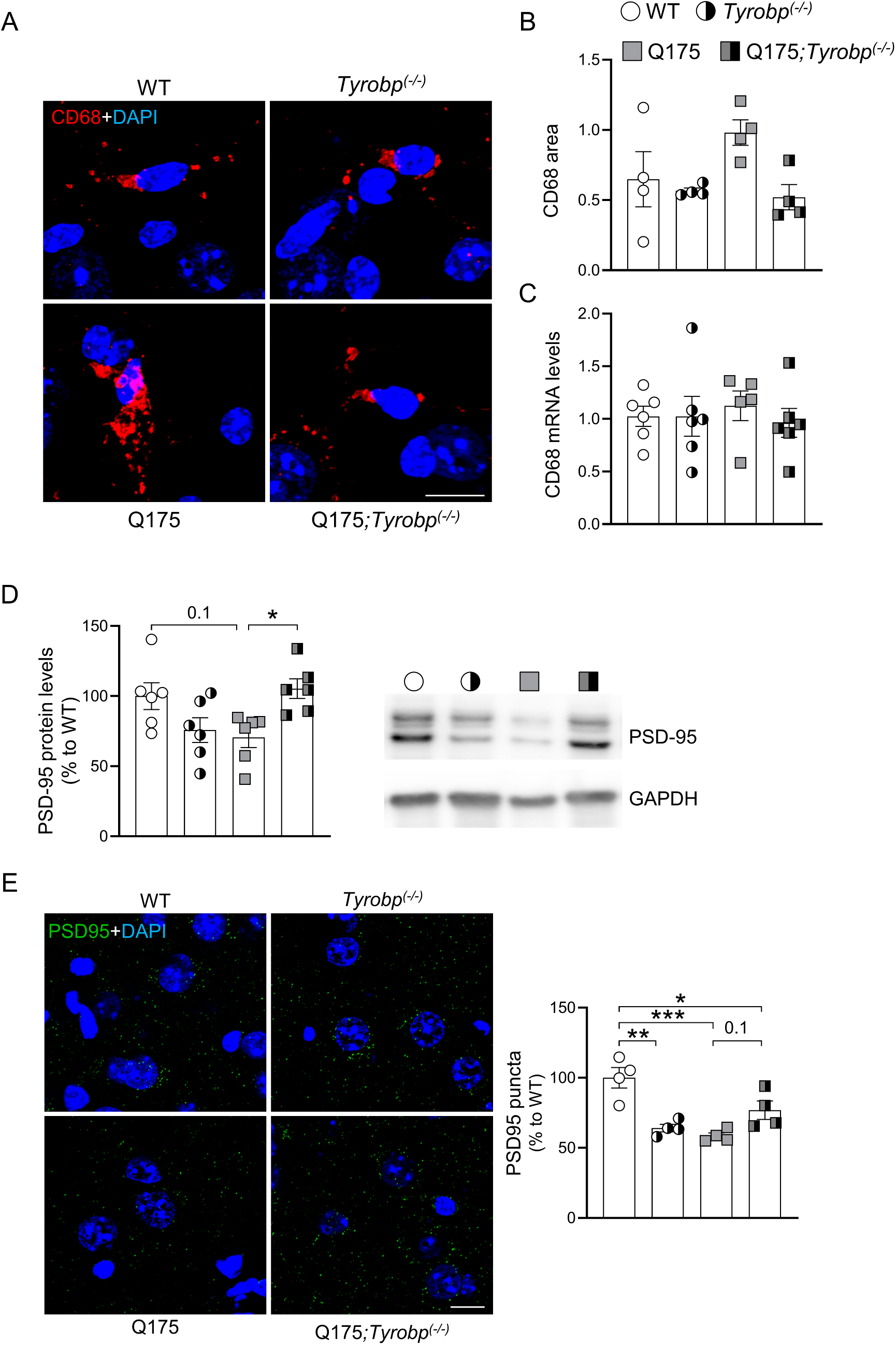
*Tyrobp* deletion prevents CD68 increase and PSD95 reduction in Q175 mice. (A) Representative images of CD68 immunostaining in the striatum from 10-month old WT and Q175 mice with (left) and without (right) *Tyrobp*. Scale bar: 10 µm. (B) Quantification of CD68-immunopositive area in the striatum (n = 4 mice per group with an average of 3 slices per mouse). (C) RT-qPCR of CD68 mRNA in the striatum in the same mice as in Fig. 2a with n = 6 mice per group. (D) Western blot and densitometric analysis of PSD95 protein in striatal samples. n = 6 mice per group. (E) Representative confocal images showing PSD95 (green) immunopositive clusters in the dorsal striatum of WT and Q175 mice with and without *Tyrobp*. Quantitative analysis is shown as mean ± SEM (n = 4 animals per group). Statistical analysis was performed using Two-Way ANOVA. *P < 0.05; **P < 0.01; ***P < 0.001. Scale bars, 10 µm.

### *Tyrobp* deletion prevents impaired motor function in 6- and 9-month-old Q175 mice

Next, we investigated whether absence of *Tyrobp* modulates motor phenotype in the Q175 mouse model. HD is characterized by decline in motor abilities, a phenotype clearly recapitulated by Q175 mice. Motor coordination was assessed by the accelerating rotarod, which also evaluates motor learning, and is highly sensitive to subtle motor changes. Balance was evaluated using the balance beam and vertical pole tests. To determine if *Tyrobp* deletion modifies the disease onset and/or improves motor behavior at symptomatic stages, motor performance was examined at two time points, 6 and 9 months of age. *Tyrobp* deletion in Q175 mice improved motor learning at both time points (Fig. 4A). Strikingly, balance deficits were detected in Q175, but not Q175;*Tyrobp*^*(-/-)*^ mice at 6 months of age (Fig. 4B,C). We also detected an overall balance improvement in Q175 *Tyrobp*^*(-/-)*^ mice at 9 months of age (Fig. 4B,C). We also noted that deletion of *Tyrobp* in the absence of HD pathology induced slight motor learning deficits, suggesting that correct *Tyrobp* gene dosage is important for striatal-dependent function. These behavioral data are compatible with a beneficial effect of the *Tyrobp* deletion on the HD phenotype.

**Fig. 4.**
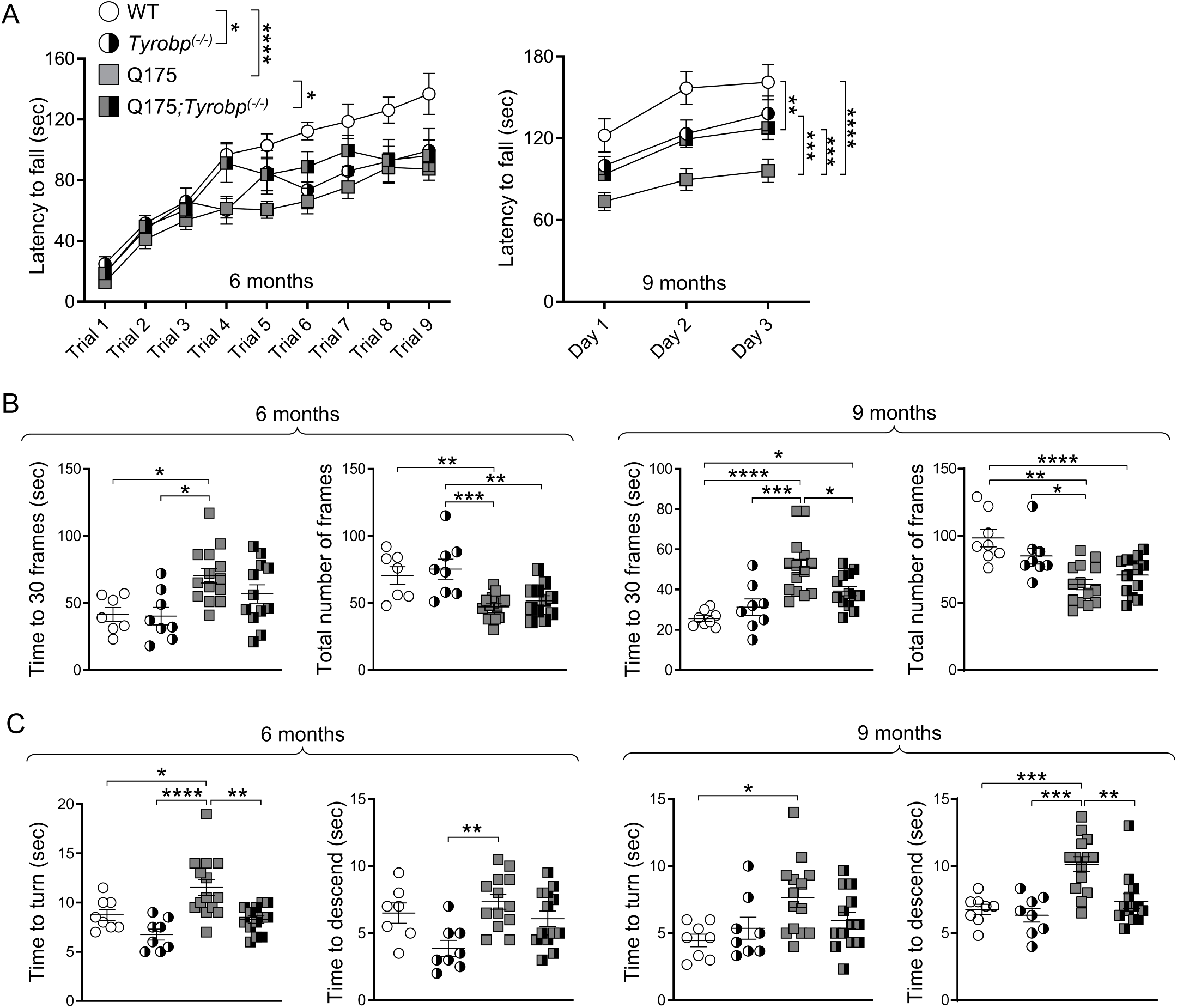
*Tyrobp* deletion ameliorates Q175 motor phenotype. Motor behavior in Q175 mice was assessed at 6 and 9 months of age. (A) Accelerating rotarod test was performed for three consecutive days (three trials per day). The latency to fall data at 6 months is represented per test and group as mean ± SEM. The latency to fall at 9 months is averaged per day (WT n = 8; *Tyrobp*^*(-/-)*^ n = 8; Q175 n = 13; Q175;*Tyrobp*^*(-/-)*^ = 12). (B) Balance beam assay is shown as time to cross 30 frames and number of frames crossed in 2 minutes (WT n = 7; *Tyrobp*^*(-/-)*^ n = 8; Q175 n = 14; Q175;*Tyrobp*^*(-/-)*^ = 13). (C) Vertical pole assay is shown as time to turn (left) and time to descend (right), which were recorded after placing the mice upwards to the pole. Three trials were conducted (WT n = 8; *Tyrobp*^*(-/-)*^; n = 8; Q175 n = 14; Q175;*Tyrobp*^*(-/-)*^ = 13). Data represent the mean ± SEM. Each point represents data from an individual mouse. Statistical analysis was performed using Two-way ANOVA followed by Bonferroni’s post hoc test, *P < 0.05; **P < 0.01; ***P < 0.001, ****P < 0.0001.

### *Tyrobp* deletion in HD mice normalizes human-specific pro-inflammatory pathways

Deletion of *Tyrobp* in AD mouse models corrects microglia-related transcriptomic abnormalities, mainly repressing the expression of inflammatory cytokines and disease-associated microglia (DAM)-related genes [21]. Here we observed that *Tyrobp* deletion in HD mice corrected microglial morphology, restored PSD-95 levels, and improved motor phenotype. We asked whether *Tyrobp* deletion would restore the overall transcriptome of HD models and/or correct pathways altered in HD human brain. We performed RNAseq analysis on striatal samples from the 10-month-old WT and Q175 mice which were tested for motor assays, including WT or homozygous KO for *Tyrobp*. Consistent with results from previous studies [18,39–45], Q175 mice showed marked transcriptomic abnormalities, with a total of 2413 differentially expressed genes (DEGs) (false discovery rate (FDR) < 0.1) relative to WT mice. Of these, 1069 were upregulated and 1344 downregulated (Fig. 5A). As expected, we detected strong downregulation of MSN identity genes, including *Drd1, Drd2, Ppp1r1b* and *Adora2a* (Supplementary Table 3). We performed gene ontology analysis using gene set enrichment analysis (GSEA). The HD transcriptome showed strong downregulation of neuronal-related genes and overall activation of development-related genes (Fig. 5B; Supplementary Table 3; Supplementary Table 4).

**Fig. 5.**
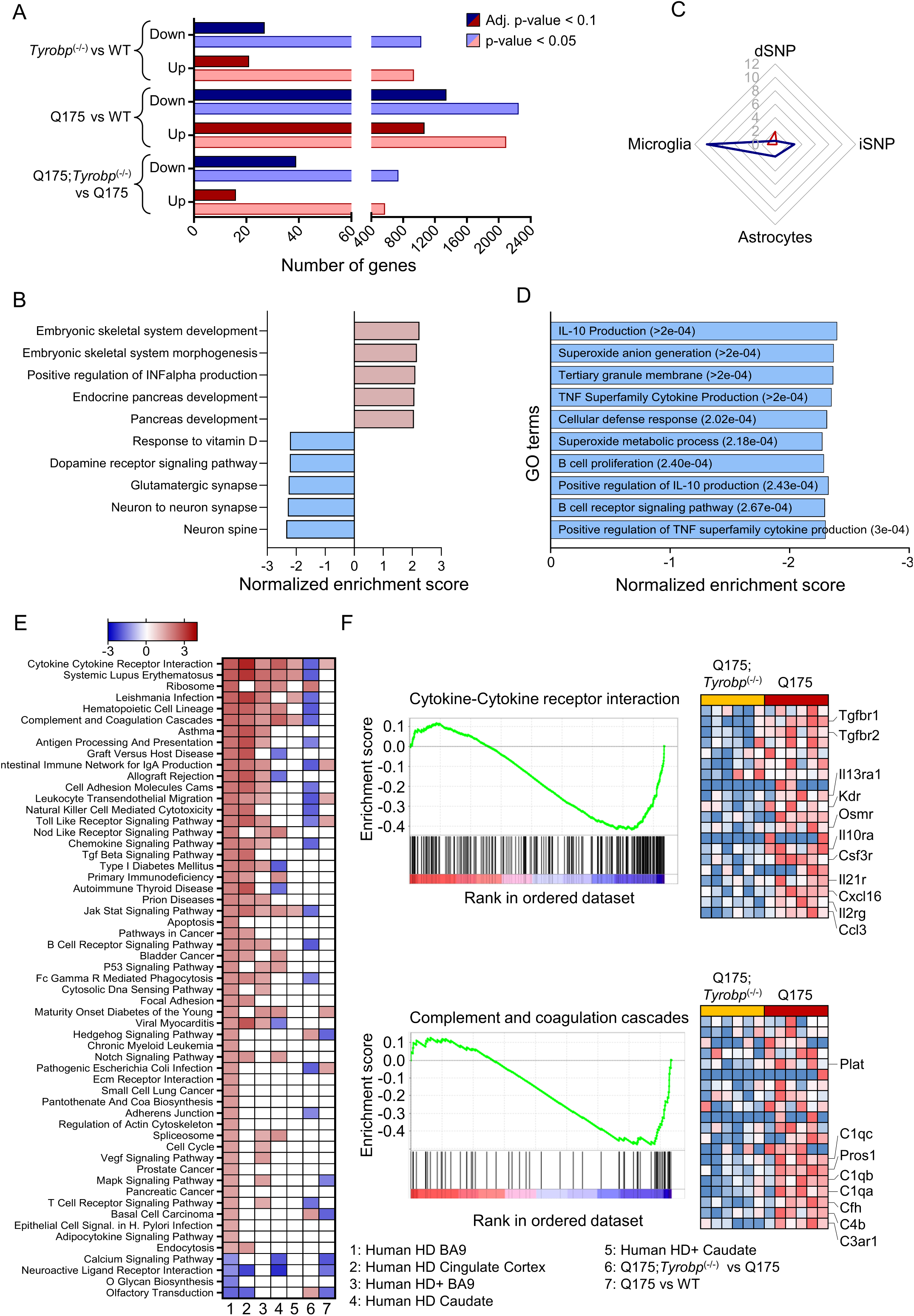
*Tyrobp* deletion reduces pathways activated in human HD brain but not in HD mouse models. Bulk RNA sequencing was performed on whole striatal RNA. (A) The number of DEGs for each data set at adjusted P < 0.1] and nominal P < 0.05 is shown. Blue bars indicate downregulated DEGs, and red bars indicate upregulated DEGs. (B) GO terms (biological process) associated with genes from Q175 vs WT data sets after GSEA analysis. (C) Cell-type enrichment analysis for Q175;*Tyrobp*^(-/-)^ vs Q175 DEGs (nominal p-value < 0.05). Lines represent –log10 p-value of Chi-square calculation of the cell type enrichment. (D) GO terms (biological process) associated to genes from Q175;*Tyrobp*^(-/-)^ vs Q175 data sets after GSEA analysis. (E) Heatmap of Normalized Enrichment Score for Kyoto Encyclopedia of Genes and Genomes (KEGG) pathways detected in human and mouse datasets. Pathways have been ranked based on descending normalized enrichment score from the Human HD BA9 dataset (Agus et al., 2019 - [15]). Column #1: Human HD Brodmann area 9 (BA9) (Agus et al., 2019 - [15]); #2: Human HD Cingulate Cortex (Al-dalahmah et al., 2020 - [70]); #3 Presymptomatic human HD BA9 vs control (Agus et al., 2019 - [15]); #4: Human HD caudate (CAU) v Control (Elorza et al., 2021 - [68]); #5 Presymptomatic human Caudate (Agus et al., 2019 - [15]); #6: Q175;*Tyrobp*^(-/-)^ vs Q175; #7: Q175 vs WT. (F) GSEA results for Cytokine-Cytokine receptor interaction and Complement and coagulation cascades pathways. Normalized gene scores are shown.

*Tyrobp* deletion in Q175 mice induced small transcriptomic changes, with 55 DEGs relative to Q175, with 16 upregulated and 39 downregulated. At a lower stringency threshold (p-value < 0.05), we detected more than 1000 DEGs (Fig. 5A; Supplementary Table 3). We confirmed the markedly decreased expression of *Tyrobp*, and importantly, that downregulated DEGs are associated with microglial cells (Fig. 5C). Again, we used GSEA to identify which biological processes and pathways were affected by *Tyrobp* deletion in Q175 mice. The most significantly downregulated biological processes were IL-10 production, superoxide anion generation, and TNF superfamily cytokine production which includes, among other genes, *Clec7a, Trem2* and *Tlr4* (Fig. 5D; Supplementary Table 4). Those biological terms are related to immune and inflammatory responses. *TREM2* and *TLR4* are increased in the putamen of HD patients and, importantly, have been identified as genetic modifiers of HD. *TREM2* R47H gene variant is associated with changes in cognitive decline in HD patients, and rs1927911 and rs10116253 *TLR4* single nucleotide polymorphisms are associated with the rate of motor decline [46]. As previously stated, genes associated with neuroinflammatory and neuroimmune responses are upregulated in the striatum and cortex of human HD brains beginning at pre-symptomatic stages [15], and these neuroinflammatory genes are unchanged from WT in the analysis of Q175 mice presented herein. To evaluate if *Tyrobp* deletion modulates pathways selectively activated in HD human brains, even if not dysregulated in the mouse, we took advantage of previously published pathway analysis performed on transcriptomic data obtained from HD human presymptomatic caudate, and symptomatic and presymptomatic cortices [15]. We also performed gene set enrichment analysis on the RNA-seq datasets used in Fig. 1 (Supplementary Table 5). The key finding is that the human pro-inflammatory gene signature is downregulated in Q175-*Tyrobp*^*(-/-)*^ mice (Fig. 5D) even though not originally increased in Q175. Pathways activated in symptomatic and presymptomatic human brains that are downregulated by *Tyrobp* deletion in Q175 brains include cytokine-cytokine receptor interaction or complement and coagulation cascades (Fig. 5E-F). Importantly, *Tyrobp* deletion in Q175 mice normalizes neither DARPP-32 levels (Supplementary Figure 2) nor the expression of MSN-specific genes (Supplementary Figure 3). These data imply that *Tyrobp* deletion does not restore neuronal pathways at the gene expression level but may be a novel target by which to reduce the expression of pro-inflammatory pathways related genes.

### Identification of *Tyrobp* as mediator of astrogliosis by integration of proteomics and transcriptomics data

Thus far, we have identified the overall transcriptomic changes induced by *Tyrobp* deletion in Q175 mice, highlighting the complement system. Next, we aimed to add another layer of analysis to identify higher-confidence candidates underlying the amelioration of Q175;*Tyrobp*^*(-/-)*^ mice motor function. To attain this, we performed proteomic analysis for striatal tissue samples from 10-month-old WT, Q175, Q175;*Tyrobp*^*(-/-)*^ and *Tyrobp*^*(-/-)*^ mice. We performed a comprehensive and quantitative workflow using data-independent acquisition (DIA) [47–49] in order to compare our groups of mice. Briefly, for quantification, all peptide samples were analyzed by DIA using variable-width isolation windows. The variable window width is adjusted according to the complexity of the typical MS1 ion current observed within certain m/z. DIA acquisitions produce complex MS/MS spectra, which are a composite of all the analytes within each selected Q1 m/z window, and subsequently, allow for highly comprehensive and accurate quantification and deep coverage. In addition, DIA workflows are not limited by the stochastic peptide MS/MS sampling biases characteristic of traditional data-dependent acquisition (DDA) mass spectrometry [49]. In order to obtain a deep spectral library for our quantitative analysis of mouse striatum from 4 different mouse strains, we first pooled aliquots of proteolytic peptides obtained after Lys-C and trypsin digestions from all samples, and further offline fractionate a subset of the pooled samples using High-pH Reversed-Phase Peptide Fractionation (HPRP). After label-free DIA acquisitions [47–49] of each of the 8 HPRP fractions and several unfractionated pooled samples using the Orbitrap Eclipse Tribrid mass spectrometer, we generated a resource of a striatum specific spectral library containing 46,185 peptides corresponding to 5,363 unique protein groups (7,950 proteins). Subsequently, each of the WT and Q175 cohort samples (with or without *Tyrobp*) was acquired in DIA mode allowing for accurate and comprehensive quantification comparing the various genotypes. Overall, from this cohort we were able to identify 3,848 quantifiable protein groups with at least 2 unique peptides. Our approach allowed us to perform in-depth proteome analysis, and covered a dynamic range over 5 orders of magnitude by abundance level (Supplementary Fig. 4A). GO biological processes and cellular compartments correlated with overall protein abundances in the anticipated manner (Supplementary Fig. 4B). Our dataset included very well-defined striatal-enriched markers, such as Ppp1r1b, Pde10a, Adcy5, and receptors, including Drd1, Adora2a and Gabra1. All protein identifications and quantifications are included in Supplementary Table 6. Partial least squares-discriminant analysis revealed highly specific and separated clustering of all genotypes, with greater separation between WT and Q175 (Fig. 6A). We detected differentially expressed proteins in all comparisons examined (DEPs, FDR < 0.05 and absolute Log_2_(fold change) > 0.2). We observed a higher number of DEPs in the Q175 vs WT comparison, in agreement with RNA-seq data, with 275 DEPs (Fig. 6B). Consensus path analysis revealed that downregulated proteins in Q175 striatum were enriched for GO terms related to the neuronal system (Fig. 6C and Supplementary Table 7). In agreement with transcriptomics data, we detected a small number of DEPs in the Q175-*Tyrobp*^*(-/-)*^ vs Q175 comparison, but importantly, these included a large reduction in C1q and GFAP protein levels when *Tyrobp* is absent (Fig. 6D). Interestingly, we detected that deletion of *Tyrobp* in Q175 increases the levels of serine protease Htra1, which is known to degrade aggregated and fibrillar tau [50]. In order to identify the most robust expression changes in Q175 and the impact of *Tyrobp* deletion in Q175 mice, we evaluated the correlation between gene and protein expression changes and examined the expression of those genes detected by both RNA-seq and proteomic profiling. Given the differences in the number of significant DEGs and DEPs, we used a different threshold for each comparison. We used genes detected as DEGs with adjusted p-value < 0.05 in Q175 vs WT comparison, and genes detected with nominal p-value < 0.05 for Q175-*Tyrobp*^*(-/-)*^ vs Q175 comparison. Remarkably, we detected high concordant changes in the proteome and transcriptome in both cases (Fig. 6E,F). We observed that *Gfap* is the gene/protein with the highest fold change. Importantly, we found that deletion of *Tyrobp* reduces both gene and protein expression of *C1q* and *Gfap*. Using immunohistofluorescence, we obtained consistent experimental evidence to support the normalization of *Gfap* expression in Q175;*Tyrobp*^*(-/-)*^ mice (Fig. 6G), which has been prioritized for validation based on the integration of RNA-seq and proteomics data.

**Fig. 6.**
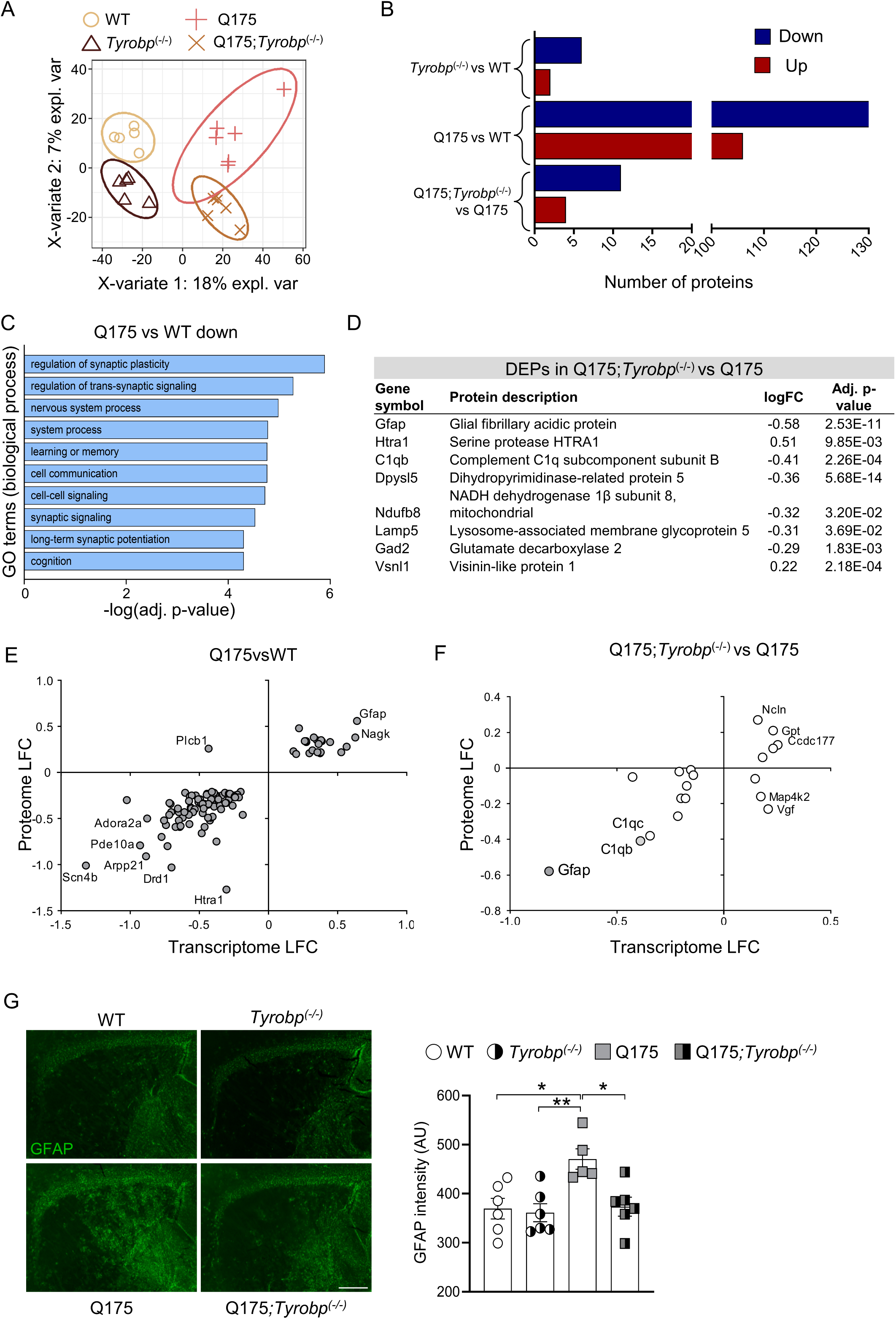
Proteomic analysis reveals reduction of astrogliosis in Q175 mice lacking *Tyrobp*. (A) Supervised Clustering using Partial Least Squares-Discriminant Analysis (PLS-DA). (B) Summary of the number of differentially expressed proteins (FDR < 0.05 and absolute Log_2_(fold change) > 0.2). Blue and red bars indicate downregulated and upregulated DEGs, respectively. (C) GO terms (biological process) associated with downregulated proteins from Q175 vs WT data sets after ConsensusPathDB over-representation analysis. (D) Top differentially expressed proteins in the striatum of Q175;*Tyrobp*^(-/-)^ vs. Q175 mice. (E,F) Plot showing Q175 vs WT (left) and Q175;*Tyrobp*^(-/-)^ vs Q175 (right) log2 fold-change (LFC) in the transcriptome and proteome. The plot only includes those genes detected as both transcripts and proteins and also differentially expressed (adj. p-value < 0.1 for Q175 vs WT comparison; and nominal p-value < 0.05 for Q175;*Tyrobp*^(-/-)^ vs Q175 comparison). Gray color denotes that the gene was differentially expressed in the transcriptome and proteome with an adj. p-value < 0.1. (G) Representative images showing GFAP (green) staining in the striatum of WT and Q175 mice with and without *Tyrobp*. Quantification of the intensity is shown as mean ± SEM (n = 5-6 mice per group). Each point represents data from an individual mouse. Statistical analysis was performed using Two-Way ANOVA. *P < 0.05; **P < 0.01. Scale bar, 500 µm.

### *Tyrobp* deletion, complement, and C1q

The integration of transcriptomic with proteomic data showed that *Tyrobp* deletion in Q175 mice reduced the expression of several genes included in complement and coagulation cascades. Also, we previously linked *Tyrobp* deficiency with downregulation of *C1q* expression in AD models [20,21]. Importantly, postmortem studies of HD human tissue have identified increases in complement components, including C1q, C3, C4, iC3b, and C9 [51]. We assessed the expression of *C1q* by qPCR and Western blot at 10-month-old in our four groups of mice. In agreement with the RNA-seq data, *C1q* mRNA and protein levels were reduced in Q175-*Tyrobp*^*(-/-)*^ mice (Fig. 7).

**Fig. 7.**
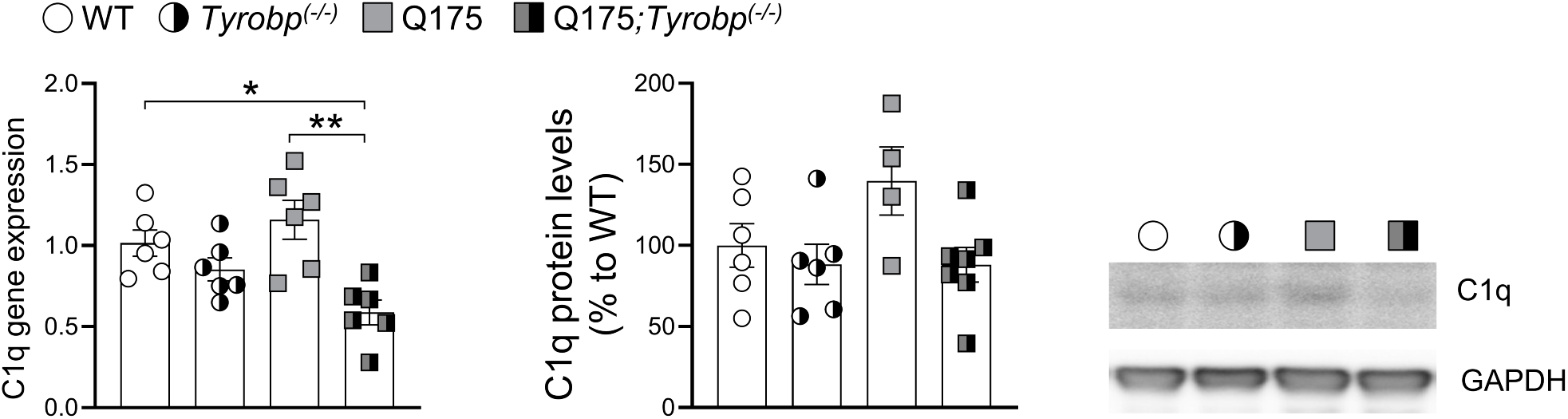
*Tyrobp* deletion reduces C1q levels in Q175 mice. (A) RT-qPCR of *C1q* mRNA in the striatum of WT and Q175 mice with and without *Tyrobp* (10 months of age), n = 6 mice per group. (B) Western blot and densitometric analysis of C1q protein in striatal samples. n = 4-6 mice per group. Quantitative analysis is shown as mean ± SEM. Each point represents data from an individual mouse. Statistical analysis was performed using Two-Way ANOVA. *P < 0.05; **P < 0.01.

### HD microglia gene dysregulation is not corrected by *Tyrobp* deletion

Cell autonomous and non-cell autonomous mechanisms may contribute to reactive microgliosis in HD brains. It has been previously demonstrated that mHtt expression in microglial cells is sufficient to elicit cell-autonomous transcriptional activation of pro-inflammatory genes in vitro [19]. However, we and others failed to detect activation of pro-inflammatory transcriptional pathways in HD mouse models. These data suggest that bulk transcriptomic analyses may not be sufficiently sensitive to detect microglial transcriptional alterations. To obtain a comprehensive and integrative picture of the transcriptional landscape of striatal HD microglia, and to evaluate its modulation by *Tyrobp* deletion, we isolated microglia from the striata of adult Q175 mice on a wild-type or homozygous *Tyrobp* KO background. Normalized counts from all samples were compared to striatal cell-type specific genes [52]. Microglial-specific genes are highly enriched in this data set compared with markers for D1- and D2-MSNs and astrocytes (Fig. 8A). Differential expression analysis comparing Q175 versus WT microglia identified mostly upregulated genes, with 228 upregulated and 4 downregulated (Fig. 8B; Supplementary Table 8). Surprisingly, pathway analysis revealed several families specifically associated with neuronal functions, including transmission across chemical synapses or neuronal system (Fig. 8D). We observed that HD microglia show higher expression of GABA-related genes (i.e., *Gabra2, Gabrb1* and *Gabrg1*) and significant enrichment for glutathione conjugation and biological oxidations, pathways that include genes related to detoxification mechanisms, e.g. *Gstm1, Gsta4* and *Gstm5*.

**Fig. 8.**
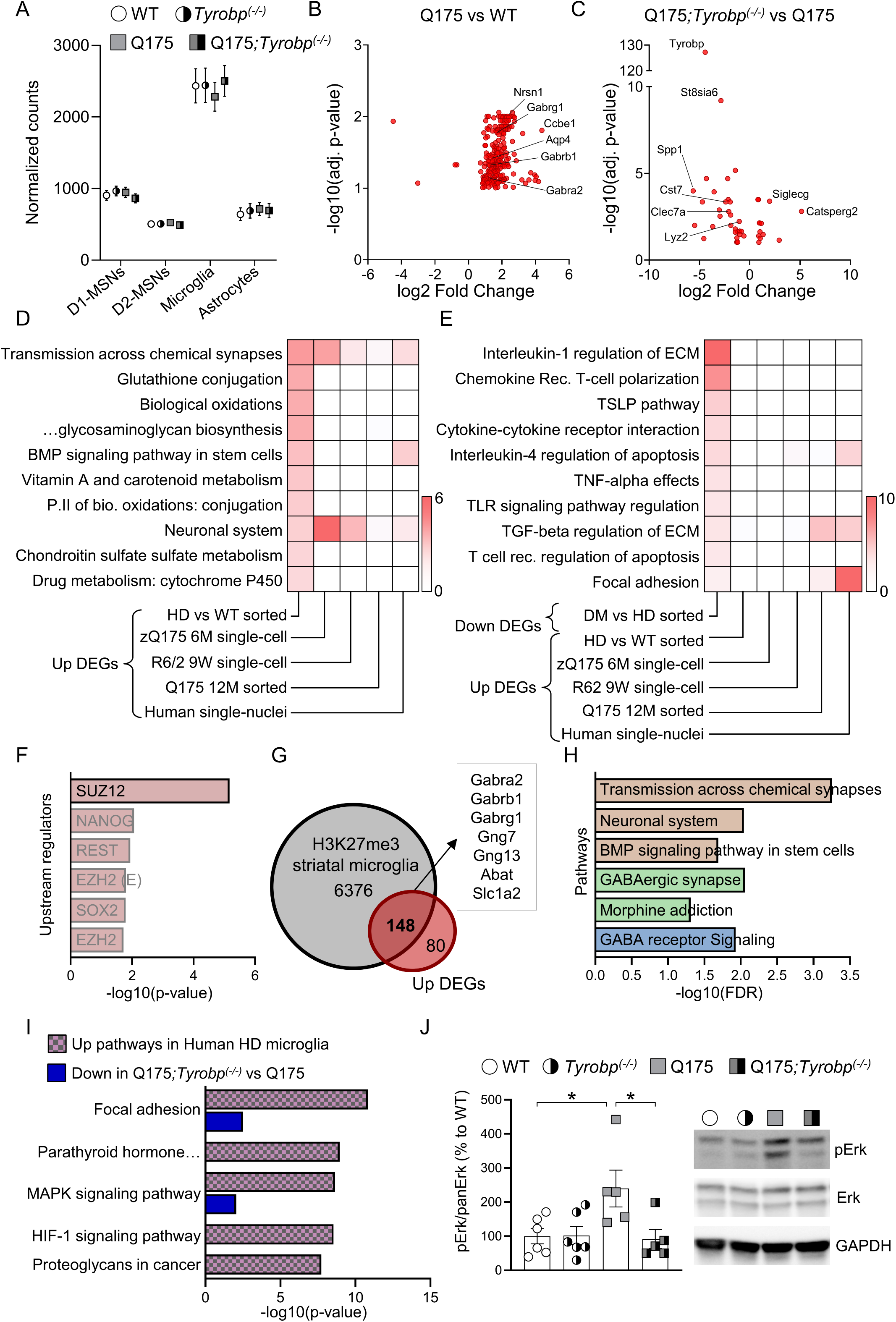
Analysis of the HD microglia-specific transcriptome and the consequences of *Tyrobp* deletion. RNA sequencing was performed on RNA purified from freshly isolated striatal microglia. (A) Analysis of global levels of cell type-specific transcripts from sorted mouse microglia RNA-seq samples using geometric mean of the normalized counts of cell type marker genes of D1-MSNs, D2-MSNs, astrocytes and microglia. (B, C) Volcano plot illustrating the DEGs identified in (B) Q175 vs WT comparison and (C) Q175;*Tyrobp*^(-/-)^ vs Q175 comparison. Only the genes with FDR < 0.1 are shown. (D) Heatmap of binomial P values for Bioplanet pathways upregulated in human and mouse datasets. (E) Heatmap of binomial P-values for Bioplanet pathways decreased in Q175;*Tyrobp*^(-/-)^, and integration with increased pathways from specified human and mouse datasets. (F) Predicted upstream regulators of upregulated genes from Q175 microglia. (G) Intersection between genes with H3K27me3 marks from striatal microglia and upregulated genes from Q175 microglia. (H) Pathways associated with upregulated DEGs with a H3K27me3 mark. Color denotes database (Brown: Bioplanet; Green: KEGG pathways; Blue: Wikpathways). (I) KEGG pathways enriched for upregulated genes in the microglial cluster of HD human snRNA-seq (pink/gray), integrated with the pathways downregulated by *Tyrobp* deletion in Q175 mice (blue). (J) Western blot and densitometric analysis of phosphorylated and total Erk protein in striatal samples. n = 5-6 mice per group. Quantitative analysis is shown as mean ± SEM. Each point represents data from an individual mouse. Statistical analysis was performed using Two-Way ANOVA. *P < 0.05.

To determine the reproducibility of the genes and pathways altered in HD microglia, we performed pathway enrichment analysis for upregulated genes identified comparing control and HD microglial clusters from publicly available single-nuclei RNA-seq experiments. These sets include data from human caudate/putamen, and zQ175 (6-month-old) and R6/2 (9-week-old) mouse models [53] and a recent study which profiled freshly isolated microglia from Q175 at 12 months of age [54]. We unexpectedly found that neuronal-related pathways are upregulated across all sets, including HD human microglia (Fig. 8D, Supplementary Table 9). Of note, pathways are shared despite the low overlap of specific genes across datasets. To identify potential key drivers of this transcriptional dysregulation, we performed ChEA analysis for upregulated genes. We observed that Suz12 is the only significant predicted upstream regulator of the up-regulated DEGs (Fig. 8F). Suz12 is a core component of the polycomb repressive complex 2 (PRC2) and is altered in HD [55]. PRC2 catalyzes histone Histone 3 Lysine 27 di- and trimethylation (H3K27me2 and H3K27me3), leading to gene repression [56]. For this reason, we intersected our upregulated DEGs with striatal microglia specific H3K27me3 marks [57], and found that 148 of the 228 up-regulated DEGs have a H3K27me3 histone mark in striatal microglia (Fig. 8G). Importantly, we observed that those genes are related to neuronal function (Fig. 8H). These data suggest that mHtt may disturb PRC2 function and thereby derepress the expression of neuronal genes in microglial cells. Whether the expression of neuronal genes in HD microglia contributes to HD pathophysiology is still unknown.

Next, we focused on how *Tyrobp* modulates the HD/Q175 microglia transcriptome. Comparing Q175;*Tyrobp*^*(-/-)*^ versus Q175 mice, we identified 43 DEGs (Fig. 8B), which primarily represented a decrease in the transcription of genes related to the *Tyrobp* network, including *Clec7a, Lyz2*, and *Spp1*. We next performed pathway enrichment analysis on the list of downregulated genes in Q175 *Tyrobp*^*(-/-)*^ versus Q175 mice (FDR < 0.1). Interleukin-1 regulation of extracellular matrix, Chemokine receptor T-cell polarization, TSLP pathway, Cytokine-cytokine receptor interaction and IL-4 regulation of apoptosis were the most significantly dysregulated pathways in the Q175 *Tyrobp*^*(-/-)*^ versus Q175 mice comparison (Fig. 8E). As we previously detected in our bulk sequencing analysis, the pathways affected by *Tyrobp* deletion in Q175 mice are not activated in HD mouse models. However, IL-4 regulation of apoptosis, TGFβ regulation of extracellular matrix and Focal adhesion are pathways downregulated by *Tyrobp* deletion in Q175 mice, that are activated in human HD microglia (Fig. 8E; Supplementary Table 10). For this reason, we next focused on microglial-specific pathways activated in HD human brains (Fig. 8I). We observed that the top pathways enriched for upregulated DEGs detected in microglial cells from HD brains were shared with those pathways downregulated in Q175:*Tyrobp*^*(-/-)*^ when compared to Q175 (Fig. 8I). IL-2 signaling pathway and focal adhesion, two upregulated Bioplanet pathways in human HD microglia were downregulated by *Tyrobp* deletion in HD mice (Fig. 8I, Supplementary Table 10).

We also noted that the MAPK signaling pathway is activated in human HD microglia and downregulated in Q175;*Tyrobp*^*(-/-)*^mice. Microglia-mediated pathogenic processes partly depend on upstream signaling events engaged by multiple pathological stimuli, probably including mHtt expression. ERK signaling regulates pro-inflammatory microglial activation in response to interferon γ [58,59]. Also, it is an upstream regulator of several DAM genes and genetic risk factors for LOAD [60]. For this reason, we evaluated Erk phosphorylation status in the striatum of our mice. Although we did not perform this analysis in a cell-type specific manner, we observed that Erk is hyperphosphorylated in the striatum of HD models (Fig. 8J), as previously described [61–63]. Importantly, *Tyrobp* deletion in Q175 mice normalized phosphorylated Erk to wild-type levels (Fig. 8J). These data demonstrate that *Tyrobp* is a candidate to not only normalize the pro-inflammatory pathways of human microglia, but also to restore, at least, Erk intracellular signaling pathways in mouse models.

## DISCUSSION

In this study, we employed multiple approaches to identify *Tyrobp* as a mediator of microglial and neuronal dysfunction in HD. We demonstrated that abrogation of endogenous *Tyrobp* in a mouse model of HD normalizes microglial morphology, impedes the reduction of PSD-95, improves motor function, and downregulates transcriptomic pathways altered in HD human brain. We extended these observations by generating and integrating transcriptomic data from HD microglia. We observed that *Tyrobp* deletion reduces the expression of genes belonging to pathways selectively activated in HD human microglia. Ultimately, our data lead to the proposed reduction of *Tyrobp* and/or its signaling pathway as a potential therapeutic approach to mitigate mHtt detrimental consequences and promote neuronal function.

*TYROBP* has been identified as a hub gene in a specific LOAD pro-inflammatory subnetwork [23]. Our group validated this observation in an APP/PSEN1 model of amyloidosis and in a model of tauopathy, where complete deletion of *Tyrobp* is phenotypically beneficial [20–22]. HD and AD share some pathological hallmarks, including altered proteostasis and detrimental pro-inflammatory responses; however, the neuroinflammatory characteristics are different. HD brains show normal levels of Ig throughout the disease course, suggesting that there is no generalized activation of the adaptive immune response [14]. In addition to AD, *TYROBP* was identified as hub gene in a microglial module conserved between human aging and neurodegenerative diseases, including PD and HD [64]. However, previous gene expression and network analyses restricted to HD samples did not identify a *TYROBP* network whose connectivity or expression is altered in HD [18,65–67]. The previous studies, however, were conducted using microarray data, since high throughput HD transcriptomic data is still scarce. Here, we used the most recent cortical and striatal RNA-seq studies [68–70] to perform pathway analysis. RNA-seq has several benefits over microarrays, including higher resolution for the identification of low-abundance transcripts and the ability to distinguish expression profiles of closely related paralogues. Differential connectivity network analysis can capture more transcriptional information between cases and controls than was previously appreciated by differential expression alone, so network-based analysis on striatal HD RNA-seq data is of great interest. Here, we observed that “TYROBP causal network in microglia” is the most enriched pathway for the upregulated genes detected across all human HD RNA-seq datasets. Also, our PPI-network results showed that *TYROBP* is a potential hub gene in HD brains.

We next provide biological validation of *Tyrobp* as a modulator of HD pathology in a mouse model. Microglial activation is a well-established hallmark of HD human brains, however, there are divergent studies regarding the microglial status in HD mouse models. We saw that striatal Q175 microglia display a pro-reactive phenotype and are increased in number, which agrees with previous studies using zQ175 mice [63], and other transgenic HD mouse models such as R6/2 [71,72]. Nonetheless, there are other studies in which these alterations are not detected [73,74]. A key observation, herein and in our previous report using a tauopathy model, is that deletion of *Tyrobp* is sufficient to normalize aberrant microglial morphology and reduce CD68 content. In accordance, we detected a preservation of PSD-95 levels in HD mice lacking *Tyrobp*. These data suggest that *TYROBP* may be a novel target to prevent synaptic dysfunction in neurodegenerative diseases. Importantly, reduction of spine density has been identified in HD human brains and across several models of HD [36–38,75,76]. In fact, excessive microglia-mediated synaptic pruning leading to synaptic dysfunction is a common pathological mechanism for many neurological and psychiatric pathologies. Although we did not directly address synaptic function in HD mice lacking *Tyrobp*, we detected an overall improvement of motor phenotype and synaptic markers, demonstrating that neuronal function is ameliorated as a consequence of *Tyrobp* deletion.

We demonstrate herein, as have many others, that HD mouse models accurately mimic human disease neuronal transcriptomic and proteomic alterations while other abnormalities, e.g. the activation of pro-inflammatory profiles, are not completely replicated. We observed that most of the inflammatory pathways transcriptionally increased or active in symptomatic and presymptomatic human caudate are downregulated by *Tyrobp* deletion in Q175 mice, despite the fact that they are not upregulated in Q175 only. We report that complement signature is downregulated in Q175;*Tyrobp*^*(-/-)*^ mice, which is in line with our previous findings in AD-related models. Although we did not detect activation of the complement pathway in HD mouse striatum, co-cultures of wild-type microglia with striatal neurons expressing mHtt displayed increased proliferation, elevated levels of cytokine IL-6 and complement components C1qa and C1qb, and take on a more amoeboid morphology [77]. These data point to a key role of *Tyrobp* in regulation of the complement system in yet another neurodegenerative disease. The integration of proteomic and transcriptomic data revealed strong overall agreement between changes in mRNA and protein levels for both induced and repressed protein/genes in response to the presence of mHtt. This concordance has also been detected for the genes altered due to the absence of *Tyrobp*.

We also detected astrogliosis in the striatum of symptomatic Q175 mice, in agreement with previous studies [74]. Importantly, we observed that *Tyrobp* deletion reduces *Gfap* mRNA and protein levels, similarly to C1q. These data agree with previous studies showing that correction of aberrant microglia impacts astrocyte activation. It was previously shown that activated microglia secrete factors, including C1q, that induce a subtype of reactive astrocytes, which are termed A1 [78]. Importantly, A1 astrocytes are abundant in HD human brain [78]. However, based on our transcriptomic data, we did not detect an A1-like gene signature in Q175 mice. In fact, little or no evidence of neurotoxic A1 astrocyte signature has been detected in Q175 mouse model [74]. Again, we contemplate a plausible scenario where deletion of *Tyrobp* has an impact on human-specific altered pathways not measured by bulk transcriptomics in HD mouse models.

We also generated an unbiased transcriptomic dataset of freshly isolated microglia from Q175 striatum, with and without *Tyrobp*, where we observed similar effects of *Tyrobp* deletion. We detected increased expression of genes related to pro-inflammatory processes such as *Pla2g5, Trcp6* or *Pdpn*. These genes have been linked to the regulation of inflammatory molecule synthesis and microglial mobility and phagocytosis [79–81]. We also report activation of glutathione conjugation genes, which may reflect a compensatory mechanism to enhance detoxification mechanisms, but also could be part of a pathological mechanism. For example, *Gstm1*, which is upregulated in HD microglia, is a glutathione S-transferase that contributes to astrocyte-mediated enhancement of microglia activation during brain inflammation [82]. Nonetheless, the overall transcriptome did not reflect immune activation.

The up-regulation of neuronal-related genes in Q175 striatal microglia, including genes encoding for GABA receptors, was unexpected. The overexpression of neuronal-related genes in HD microglia was present in other HD models and also in single-nuclei from HD human cortex. A potential mechanism is that perhaps mHtt disrupts the PRC2 complex, causing a de-repression of neuronal genes in microglia. Recent studies reported expression of GABA receptors in microglia [83–85], which are essential for inhibitory connectivity. Removal of microglial GABA receptors alters inhibitory connectivity, induces behavioral abnormalities in mice and, importantly, causes a downregulation of synaptic pruning-related genes [85]. Although the consequences of GABA receptor overexpression in microglia are not known, one plausible consideration is that the overexpression of GABA receptors in HD microglia is a compensatory mechanism to attenuate mHtt-induced pro-inflammatory responses. *In vitro*, GABA reduces LPS-induced microglial IFN-γ, IL-6 and TNFα production [83]. GABA receptor protein levels are increased in a facial nerve axotomy model, where activation of GABA B receptors, again, strongly decreases LPS-induced secretion of certain cytokines, including IL-6 and IL-12p40 [84]. These data suggest that HD microglia overexpress GABA receptor in an attempt to reduce striatal inflammation as one of several potentially compensatory mechanisms activated in the striatum of HD models [86]. Here, we detected overexpression of *Fbxo2* and *Scrg1* in HD microglia. *Fbxo2* mediates clearance of damaged lysosomes and modifies neurodegeneration in Nieman-Pick C models [87], suggesting that HD microglia may be attempting to improve lysophagy pathways. *Scrg1* suppresses LPS-induced Ccl22 production in monocyte/macrophage-like cells [88].

## CONCLUSIONS

In summary, we demonstrate that although microglia from HD mouse models do not fully recapitulate the human HD microglia transcriptome, deletion of *Tyrobp* in a full-length HD mouse model corrects potentially pathogenic pathways activated in HD microglia. Therefore, *Tyrobp* deletion improves the behavior of APP/PSEN1, MAPT^P301S^ and Q175 mice, in each case, decreasing many of the same genes even if they are not pathologically increased in the presence of *Tyropb*. These data imply that downregulation of this network in the presence of these proteinopathies is beneficial, regardless of baseline levels and MSN transcriptional dysregulation.

## METHODS

### Data Sources and Wikipathway enrichment analysis

To identify the pathways enriched for upregulated genes in HD human brains, we used publicly available HD human transcriptomic data providing differentially expressed genes or pathway enrichment analysis. We filtered all datasets on gene expression omnibus (GEO) or European Nucleotide Archive (ENA) databases based on the organism and considering both RNA-seq and microarray sequencing methods. The query was limited to all datasets relevant to HD human brain research and high throughput genomics experiments which would include up to 6 entries. PRJEB44140 (RNA-seq, striatum [68]), GSE64810 (RNA-seq, Brodmann Area 9, [69]), Al-Dalahmah et al., 2020 [70] (RNA-seq, anterior cingulate cortex, GSE26927 (microarray, caudate, [17]), GSE3790 (microarray, caudate, Broadmann area 4 and cerebellum, [89]) were analyzed. Pathway enrichment analysis was performed using the EnrichR package [90] considering only upregulated differentially expressed protein-coding genes (Fold Change > 0.5 and adjusted p-value < 0.05). Significant pathways were identified by Fisher’s exact test with adjusted p-value < 0.05. All gene sets provided a list of differentially expressed genes except GSE26927. For GSE26927 we used GEO2R online platform to retrieve DEGs. Data was log normalized and Benjamini & Hochberg (FDR) method was used to adjust the p-values. GSE79666 dataset, although it contains RNA-seq data of the BA4 motor cortex of 7 control and 7 Huntington’s disease patients, was excluded because only 51 DEGs were detected. For GSE3790, which contains microarray data from the caudate, cortex and cerebellum, the authors do not provide adjusted p-values. We set up the threshold as nominal p-value < 0.001; as suggested by the authors.

### Network construction of protein–protein interaction

To inquire into the role of common upregulated DEGs across HD brains, we analyzed these genes by means of STRING v11.5 tool. The minimum required interaction score was set to high confidence (0.7). Text mining, experiments, databases, co-expression, neighborhood, gene fusion and co-occurrence were used as prediction methods. The plug-in Molecular Complex Detection (MCODE), a well-known automated method to find highly interconnected subgraphs as molecular complexes or clusters in large PPI networks, was used to screen the modules or clusters of PPI network in Cytoscape.

### Mice

Animal procedures were conducted in accordance with the NIH Guidelines for the Care and Use of Experimental Animals and were approved by the Institutional Animal Care and Use Committee at Icahn School of Medicine at Mount Sinai. All mice were on a C57Bl6/J background. Heterozygote Q175 (JAX #029928) and *Tyrobp* knockout (*Tyrobp*^(**−**/**−**)^) [28] mice were obtained from Jackson Laboratories and Taconic/Merck Laboratory respectively. Mice were kept in a 12-hour light-dark cycle with ad libitum access to food and water at Icahn School of Medicine at Mount Sinai, NY, USA.

### Tissue extraction

10-months-old mice were anesthetized with pentobarbital (50 mg/kg intraperitoneal) and perfused with ice-cold phosphate-buffered saline (PBS). Brains were removed, hemispheres were sagittaly separated, and the striatum was dissected from the left hemisphere and flash frozen. The right hemisphere and hindbrain were post-fixed in 4% paraformaldehyde (PFA) for 48h and prepared for histological analyses. The frozen striatum was used for RNA extraction and RT-qPCR as well as western blot analysis.

### Immunohistofluorescence analysis

30 μm sagittal free-floating sections were washed with 1X Tris Buffered Saline (TBS), blocked at RT for 1h (5% goat serum, 0.25% Triton X-100, 1X TBS), and then incubated at 4ºC overnight (O/N) with mouse anti-DARPP-32 (1:2000, sc-271111, Santa Cruz), rabbit anti-Iba1 (1:500, 019-1974, Wako), rat anti-CD68 (1:500, MCA1957, Bio-Rad) and mouse anti-PSD-95 (1:500, MAB1596, Millipore) antibodies in 1X TBS + 0.05% Triton X-100 with 1% goat serum. Sections were washed with 1X TBS + 0.1% Triton X-100 and incubated with the appropriate secondary antibody: anti-rabbit Alexa 488 (1:400, #A-11034, Thermo Fisher Scientific), anti-rat Alexa 594 (1:500, #A-11007, Thermo Fisher Scientific). Images were obtained with Zeiss 700 confocal microscope (Zeiss, Thornwood, USA). Morphological analysis on microglia were performed as described in [91].

### RNA extraction and qPCR analysis

RNAs were isolated from mice striata using the QIAzol® Lysis Reagent (Qiagen) and the miRNeasy® Micro Kit (Qiagen). 500 ng of total RNAs were reverse transcribed using the High-Capacity RNA-to-cDNA Kit (ThermoFisher, #4387406). cDNAs were subjected to real-time qPCR in a Step-One Plus system (Applied Biosystem) using The All-in-One qPCR Mix (GeneCopoeia, #QP001-01). Sequences of oligonucleotides used are: Mm99999915_g1_Gaphd (4331182, ThermoFisher),

C1q: C1q: Fwd 5′-AACCTCGGATACCAGTCCG-3′; Rev 5′-ATGGGGCTCCAGGAAATC-3′

CD68: Fwd 5′-TGTCTGATCTTGCTAGGACCG-3′; Rev 5′-GAGAGTAACGGCCTTTTTGTGA-3′

The individual value for each test transcript, performed in triplicate, was normalized to the abundance of GAPDH. Relative quantification was performed using the ΔΔCt method [92] and was expressed as fold change relative to control by calculating 2-ΔΔCt.

### RNA-sequencing

Bulk RNA sequencing was performed by Novogene (https://en.novogene.com) using Illumina Novaseq 6000 S4 flow cells. Microglia RNA sequencing was performed by Genomics Technology Facility, Department of Genetics and Genomic Sciences, Icahn School of Medicine at Mount Sinai using SMART-Seq v4 Ultra Low Input. Samples with RNA integrity number (RIN) > 8 were used. Non-directional libraries were constructed with a NEB kit using the manufacturer’s protocol. RNA sequencing assays were performed after ribosomal RNA depletion by Ribo-Zero. FASTQ files were aligned to the annotated *Mus musculus* reference genome version GRCm38 using STAR (v2.5). FeatureCounts was used to quantify gene expression at the gene level based on UCSC gene model. Genes with at least 1 count per million in at least one sample were considered expressed and retained for further analysis. Differential expression analysis between two conditions/group was performed using the DESeq2 R package (1.14.1). The resulting P-values were adjusted using Benjamini and Hochberg’s approach for controlling the False Discovery Rate (FDR).

### KEGG Pathway and Reactome Pathway Analysis

GO terms, Bioplanet, KEGG pathway analysis of differentially expressed genes was performed using Enrichr tool [90]. Pathways with P-value < 0.05 were considered significantly enriched by differential expressed genes if they had a minimum of 3 genes associated.

### Gene set enrichment analysis

In order to understand the biology underlying the gene expression profile, we performed pathway analysis using GSEA. We used GSEA because it is a threshold-free method that can detect pathway changes more sensitively and robustly than some methods [93]. The following options were selected or input into the software for analysis: gene set database: - Canonical pathways: c2.cp.v72.symbols.gmt; GO terms: c5.go.v7.2.symbols.gmt; KEGG pathways: c2.cp.kegg.v7.4.symbols.gmt. Number of permutations: 1,000. Permutation type: gene_set. Chip platform: Mouse_ENSEMBL_Gene_ID_Human_Orthologs_ MSigDB.v7.2.chip.

### Proteomics analysis

#### Proteolytic digestion and desalting

The striatum was dissected from the brains of 4 different mouse strains with 5 or 6 replicates each: strains used and compared were wild-type (WT, n=5), *Tyrobp*^(-/-)^ (n=5), Q175 (n=6) and Q175;*Tyrobp*^(-/-)^ (n=6). Frozen mouse striatum was subjected to 400 μL of lysis buffer composed of 8 M urea, 2% SDS, 200 mM triethylammonium bicarbonate (TEAB), pH 8.5, 75 mM NaCl, 1 μM trichostatin A, 3 mM nicotinamide, and 1x protease/phosphatase inhibitor cocktail, and homogenized for 1 cycle with a Bead Beater TissueLyser II (QIAGEN, Germantown, MD) at 25 Hz for 1.5 min. Lysates were clarified by spinning at 15,700 x *g* for 15 min at 4°C, and the supernatant containing the soluble proteins was collected. Protein concentrations were determined using a Bicinchoninic Acid Protein (BCA) Assay (Thermo Fisher Scientific, Waltham, MA), and subsequently 100 μg of protein from each sample were aliquoted and samples were brought to an equal volume using water. Samples were then solubilized using 4% SDS, 50 mM TEAB at a pH ∼7.55. Proteins were reduced using 20 mM DTT in 50 mM TEAB for 10 min at 50°C followed by 10 min at RT, and protein were subsequently alkylated using 40 mM iodoacetamide in 50 mM TEAB for 30 min at RT in the dark. Samples were acidified with 12% phosphoric acid to obtain a final concentration of 1.2% phosphoric acid, and diluted with seven volumes of S-Trap buffer (90% methanol in 100 mM TEAB, pH ∼7). Samples were then loaded onto the S-Trap micro spin columns (Protifi, Farmingdale, NY), and spun at 4,000 x *g* for 10 seconds. The S-Trap columns were washed with S-Trap buffer twice at 4,000 x *g* for 10 seconds each, before adding a solution of Sequencing-Grade Endoproteinase Lys-C (Sigma, Atlanta, GA) in 50 mM TEAB at a 1:20 (w/w) enzyme:protein ratio at 37°C for 2 hours. A solution of sequencing grade trypsin (Promega, San Luis Obispo, CA) in 50 mM TEAB at a 1:25 (w/w) enzyme:protein ratio was then added, and after a 1-hour incubation at 47°C, trypsin solution was added again at the same ratio, and proteins were digested overnight at 37°C.

Peptides were sequentially eluted with 50 mM TEAB (spinning for 1 min at 1,000 x *g*), 0.5% formic acid (FA) in water (spinning for 1 min at 1,000 x *g*), and 50% acetonitrile (ACN) in 0.5% FA (spinning for 1 min at 4,000 x *g*). After vacuum drying, samples were resuspended in 0.2% FA in water, desalted with Oasis 10-mg Sorbent Cartridges (Waters, Milford, MA). All samples were vacuum dried and resuspended in 0.2% FA in water at a final concentration of 1 μg/μL. Finally, indexed retention time standard peptides (iRT; Biognosys, Schlieren, Switzerland) [94] were spiked in the samples according to manufacturer’s instructions.

#### Generation of the spectral library

For generating the spectral library, 15 μL from each digested striatum sample were pooled, and subsequently, an aliquot of 100 μg digested protein (from the pooled mouse striatum) was vacuum dried, resuspended in 300 μL of 0.1% trifluoroacetic acid (TFA), and fractionated using the Pierce High-pH Reversed-Phase Peptide Fractionation (HPRP) Kit (Thermo Fisher Scientific, Rockford, IL) according to manufacturer’s instructions. Eight fractions were eluted using 5%, 7.5%, 10%, 12.5%, 15%, 17.5%, 20% and 50% ACN in 0.1% triethylamine, then the 8 collected fractions were vacuum dried, and each fraction was resuspended in 12.5 μL of 0.2% FA. iRT peptides were spiked in the samples according to manufacturer’s instructions. To build the spectral library each fraction was acquired by DIA by LC-MS/MS (see below). Additionally, triplicate DIA measurements of the unfractionated pooled samples were performed.

#### Mass spectrometric analysis: Data-independent acquisition (DIA)

LC-MS/MS analyses were performed on a Dionex UltiMate 3000 system online coupled to an Orbitrap Eclipse Tribrid mass spectrometer (Thermo Fisher Scientific, San Jose, CA). The solvent system consisted of 2% ACN, 0.1% FA in water (solvent A) and 98% ACN, 0.1% FA in water (solvent B). Briefly, proteolytic peptides were loaded onto an Acclaim PepMap 100 C18 trap column with a size of 0.1 × 20 mm and 5 µm particle size (Thermo Fisher Scientific) for 5 min at 5 µL/min with 100% solvent A, loading an amount of 800 ng for each of the cohort samples and the unfractionated pool samples, and 400 ng for the HPRP fractions. Peptides were eluted on an Acclaim PepMap 100 C18 analytical column sized as follows: 75 µm x 50 cm, 3 µm particle size (Thermo Fisher Scientific) at 0.3 µL/min using the following gradient of solvent B: 2% for 5 min, linear from 2% to 25% over 96 min, linear from 25% to 40% over 23 min, linear from 40% to 50% over 6 min, and up to 80% in 1 min with a total gradient length of 170 min.

All samples - for the generation of the spectral library and for the final quantitative analysis of the cohort samples - were analyzed in DIA mode. Full MS spectra were collected at 120,000 resolution (AGC target: 3e6 ions, maximum injection time: 60 ms, 350-1,650 m/z), and MS2 spectra at 30,000 resolution (AGC target: 3e6 ions, maximum injection time: Auto, NCE: 27, fixed first mass 200 m/z). The DIA precursor ion isolation scheme consisted of 26 variable windows covering the 350-1,650 m/z mass range with an overlap of 1 m/z (Supplementary Table 11) [95].

### Data analysis

#### Spectral library generation

The DIA spectral library was generated directly in Spectronaut (version 15.1.210713.50606; Biognosys) using Biognosys (BGS) default settings and a mouse UniProtKB-TrEMBL database (86,521 entries, release 08/2021). Briefly, for the Pulsar search, trypsin/P was set as the digestion enzyme and 2 missed cleavages were allowed. Cysteine carbamidomethylation was set as fixed modification, and methionine oxidation and protein N-terminus acetylation as variable modifications. Identifications were validated using 1% false discovery rate (FDR) at the peptide spectrum match (PSM), peptide and protein levels, and the best 3-6 fragments per peptide were kept. The final spectral library contains 46,185 modified peptides and 5,363 protein groups, and can be found in Supplementary Table 6.

#### DIA data processing and statistical analysis

DIA data was processed in Spectronaut (version 15.1.210713.50606) using the previously described library. Data extraction parameters were set as dynamic and non-linear iRT calibration with precision iRT was selected. Identification was performed using a precursor PEP cut-off of 0.2 and a 1% precursor and protein q-value (experiment). Quantification was based on MS2 area, and local normalization was applied.

Differential protein expression analysis was performed using paired t-test, and p-values were corrected for multiple testing, specifically applying group wise testing corrections using the Storey method [96]. Protein groups with at least two unique peptides, q-value < 0.05, and absolute Log2(fold-change) > 0.2 were considered to be differentially expressed (Supplementary Table 6).

#### Statistical Processing

Partial least squares-discriminant analysis (PLS-DA) of the proteomics data was performed using the package mixOmics [97] in R (version 4.0.2; RStudio, version 1.3.1093).

#### Pathway Analysis

An over-representation analysis (ORA) was performed using Consensus Path DB-mouse (Release MM11, 14.10.2021) [98,99] to evaluate which gene ontology terms were significantly enriched. Gene ontology terms identified from the ORA were subjected to the following filters: q-value < 0.05, term category = b (biological process), and term level > 1.

#### Microglia isolation

##### Tissue Dissociation

Striatal cell isolation was performed as previously described [100]. Briefly, animals were deeply anesthetized with an intraperitoneal injection of pentobarbital (50 mg/kg IP) and transcardially perfused with 15 ml ice-cold, calcium- and magnesium-free phosphate-buffered saline (PBS, pH 7.3-7.4) The brains were quickly removed and striata were dissected on a cooled petri dish and placed in ice-cold Hibernate-A medium. Striata from each mouse were gently dissociated mechanically in Hibernate-A medium in a 1 ml Dounce homogenizer using the loose pestle. The homogenized tissue was then sieved through a 70μm cell strainer and transferred to a 15 ml falcon tube. The homogenates were pelleted at 450xg for 6 min at 4°C. Pellets were resuspended in PBS to a final volume of 1.5 ml. 500 μL of freshly prepared isotonic percoll solution (pH7.4) was then added to each sample (final volume: 2 ml) and mixed well. Percoll was rendered isotonic by mixing 1 part of 10x PBS (pH 7.3-7.4) with 9-parts of percoll. The percoll solution was mixed properly with the cell suspension, after which 2 ml of PBS were gently layered on top of it creating two separate layers. The samples were centrifuged for 10 min at 3000xg. The upper layers were aspirated, leaving about 500 μL as some cells float in percoll just above the pellet. The cells were then washed once in PBS making sure not to resuspend the pellet. This was achieved by gently adding 4 ml PBS, closing the tube, and holding it in a horizontal position, and gently tilting it 145 degrees in order to mix the remaining percoll with the added PBS. The cells were then pelleted by centrifuging them at 450xg for 10 min at 4°C. The resulting toal striatal cell pellet was used for the magnetic sorting of microglia as described below.

#### Microglia Isolation

Microglia were isolated from 1 striatum/sample via magnetic-activated cell sorting (MACS) using mouse anti-CD11b (for microglia) paramagnetic nanobeads (Miltenyi) according to the manufacturer’s instructions with some modifications. The MACS buffer used consisted of 1.5% bovine serum albumin (BSA) diluted in PBS from a commercial 7.5% cell-culture grade BSA stock (Thermo Fisher Scientific). For the microglial isolation, total striatal cell pellets after percoll (see above) were re-suspended in 90 μL MACS buffer and 10 CD11b beads (Miltenyi). Cells were then incubated for 15 min at 4°C. Excess beads were washed with 1 ml MACS buffer and the cells pelleted at 300 rcf for 5 min at 4°C. The cells were then passed through an MS MACS column attached to a magnet whereby CD11b positive cells stay attached to the column, whereas unlabeled cells flowed through the column. After washing the columns three times with MACS buffer, microglia were flushed from the column with 1 ml MACS buffer and pelleted at 300 rcf for 5 min at 4°C. Cell pellets were lysed in QIAzol (Qiagen product code 79306), snap-frozen in dry ice and stored at -80°C until RNA extraction.

#### Western blot

Flash frozen striatal samples were homogenized in a RIPA buffer (Pierce; 89900) containing freshly added phosphatase (Pierce) and protease (Roche) inhibitors, centrifuged for 20 min at 15,000 g and the supernatant was collected. Protein concentration was determined using the BCA method. For each sample, 20 μg of protein was resolved in 4%-12% Bis/Tris-acrylamide gradient gels (BioRad) and transferred to nitrocellulose membranes. Membranes were incubated with the following primary antibodies: anti-PSD-95 clone 6G6-1C9 (1:1000, MAB1596, Millipore), anti-phospho-p44/42 MAPK (Erk1/2) (Thr202/Tyr204) (1:1000, #9101, Cell Signaling), anti-p44/42 MAPK (Erk1/2) Antibody (1:1000, #9102, Cell Signaling), anti-C1q [4.8] (1:500, ab182451, Abacam), anti-DARPP-32 (1:1000, #2306, Cell Signaling), anti-Iba1 (1:1000, 016-20001, Wako) and anti-GAPDH (D16H11) (1:2000, #5174, Cell Signaling) antibodies. The secondary HRP conjugated antibodies included anti-rabbit (1:2000, PI-1000, Vector laboratories) and anti-mouse (1:2000, PI-2000, Vector laboratories). Following development with ECL (Pierce®) pictures was acquired using a Fujifilm ImageReader LAS-4000, and bands were quantified using ImageJ (Fiji software package). Protein levels were normalized to GAPDH, and mean values (Ns specified in the figure legend) were normalized to Control = 100.

#### Behavior

##### Accelerating rotarod

Motor learning was evaluated as in [101,102]. With neither training nor habituation sessions, mice were placed on a motorized rod (3 cm diameter) with a gradual speed increase from 4 to 40 RPMs for 5 min. Each mouse was evaluated over three days, conducting three trials per day, with an intertrial interval of 1 h. Latency to fall was recorded. Trials were included only if the mouse placed all four paws on the rod.

##### Balance beam test

Balance was evaluated as in [101,102]. The beam consisted of an 85 cm long wooden prism, divided into 5 cm frames, with a 1 cm face, placed 40 cm above the bench surface. The test consisted of two sessions, the training and the testing, separated by 4 h. In both training and testing sessions mice walked along the beam for 2 min. During the testing session latency to cover 30 frames and total distance traveled were measured.

##### Vertical pole test

Balance was evaluated as in [101,102]. The pole test was consisted of a 60 cm wooden cylinder (1 cm diameter) wrapped in tape to facilitate walking. Mice were trained for two consecutive days and tested on the third day. Three trials per session were conducted. Mice were placed with heads facing upward just below the top of the pole. Both time to complete a turn, in other words, orient the body downward, and time to climb down (time to descend) the pole were measured. Trials in which mice descended the pole without complete reorientation were counted as error trials and analyzed separately.

#### Statistical analysis

The non-genomic data (Figs. 2, 3, 4, 6G, 7, 8J) were analyzed with GraphPad Prism 8. Graphs represent the mean of all samples in each group ± SEM. Sample sizes (n values) and statistical tests are indicated in the figure legends. A two-way ANOVA followed by a Bonferroni’s post-hoc test was used for multiple comparisons. A Student’s t-test was used for simple comparisons. Significance is reported at *p < 0.05, **p < 0.01, ***p < 0.001.

## ABBREVIATIONS

HD: Huntington’s disease
HTT: Huntingtin human gene
MSNs: Medium spiny neurons
Htt: Huntingtin mouse protein
mHtt: mutant huntingtin
Tyrobp: TYRO protein tyrosine kinase-binding protein
AD: Alzheimer’s disease
LOAD: Late onset Alzheimer’s disease
DEGs: Differentially expressed genes
PSD-95: Post-synaptic density 95
DAM: Disease-associated microglia
GSEA: Gene set enrichment analysis
DIA: Data-independent acquisition
FDR: False discovery rate
DEPs: Differentially expressed proteins
PRC2: Polycomb repressive complex 2
H3K27me2: Histone 3 Lysine 27 dimethylation
H3K27me3: Histone 3 Lysine 27 trimethylation
Erk: Extracellular signal-regulated kinase
Iba1: Ionized calcium binding adaptor molecule 1
RNAseq: Ribonucleic acid sequencing
qPCR: Quantitative polymerase chain reaction
SEM: Standard error of the mean
TBS: Tris-buffered saline
PBS: Phosphate-buffered saline
WT: wild-type
Gfap: Glial fibrillary acidic protein
IL-: Interleukin-
PPI: Protein-Protein interaction
MCODE: Molecular Complex Detection

## AUTHOR CONTRIBUTIONS

JCM and MEE designed the study. JCM, DM, AR, BWH, SC performed and analyzed the experiments. JB, BS and LE performed and analyzed the proteomics experiments. JCM and MEE wrote the manuscript. All authors read and approved the final manuscript.

## AVAILABILITY OF THE DATA

Raw data and processed information of the RNA sequencing experiments generated in this article were deposited at the Gene Expression Omnibus repository under the accession numbers GSE193573 (bulk RNA sequencing) and GSE195633 (microglia RNA sequencing). Raw data and complete MS data sets have been uploaded to the Center for Computational Mass Spectrometry, to the MassIVE repository at UCSD, and can be downloaded using the following link: https://massive.ucsd.edu/ProteoSAFe/dataset.jsp?task=2194e8c2d8cd479780e75e917f946e2d (MassIVE ID number: MSV000088643; ProteomeXchange ID: PXD030747).

## REFERENCES

1. HDCRG. A novel gene containing a trinucleotide repeat that is expanded and unstable on Huntington’s disease chromosomes. The Huntington’s Disease Collaborative Research Group. Cell. United States; 1993;72:971–83.

2. Kassubek J, Bernhard Landwehrmeyer G, Ecker D, Juengling FD, Muche R, Schuller S, et al. Global cerebral atrophy in early stages of Huntington’s disease: quantitative MRI study. Neuroreport. England; 2004;15:363–5.

3. Vonsattel JP, DiFiglia M. Huntington disease. J Neuropathol Exp Neurol. England; 1998;57:369–84.

4. Han I, You Y, Kordower JH, Brady ST, Morfini GA. Differential vulnerability of neurons in Huntington’s disease: The role of cell type-specific features. J Neurochem. 2010;113:1073–91.

5. Rosas HD, Koroshetz WJ, Chen YI, Skeuse C, Vangel M, Cudkowicz ME, et al. Evidence for more widespread cerebral pathology in early HD: an MRI-based morphometric analysis. Neurology. United States; 2003;60:1615–20.

6. Creus-Muncunill J, Ehrlich ME. Cell-Autonomous and Non-cell-Autonomous Pathogenic Mechanisms in Huntington’s Disease: Insights from In Vitro and In Vivo Models. Neurother J Am Soc Exp Neurother. 2019;16:957–78.

7. Yang H-M, Yang S, Huang S-S, Tang B-S, Guo J-F. Microglial Activation in the Pathogenesis of Huntington’s Disease. Front Aging Neurosci. Switzerland; 2017;9:193.

8. Ferrante RJ, Gutekunst C a, Persichetti F, McNeil SM, Kowall NW, Gusella JF, et al. Heterogeneous topographic and cellular distribution of huntingtin expression in the normal human neostriatum. J Neurosci. 1997;17:3052–63.

9. Pavese N, Gerhard A, Tai YF, Ho AK, Turkheimer F, Barker RA, et al. Microglial activation correlates with severity in Huntington disease: a clinical and PET study. Neurology. United States; 2006;66:1638–43.

10. Tai YF, Pavese N, Gerhard A, Tabrizi SJ, Barker RA, Brooks DJ, et al. Microglial activation in presymptomatic Huntington’s disease gene carriers. Brain. England; 2007;130:1759–66.

11. Sapp E, Kegel KB, Aronin N, Hashikawa T, Uchiyama Y, Tohyama K, et al. Early and progressive accumulation of reactive microglia in the Huntington disease brain. J Neuropathol Exp Neurol. England; 2001;60:161–72.

12. Politis M, Lahiri N, Niccolini F, Su P, Wu K, Giannetti P, et al. Increased central microglial activation associated with peripheral cytokine levels in premanifest Huntington’s disease gene carriers. Neurobiol Dis. United States; 2015;83:115–21.

13. Silvestroni A, Faull RLM, Strand AD, Moller T. Distinct neuroinflammatory profile in post-mortem human Huntington’s disease. Neuroreport. England; 2009;20:1098–103.

14. Bjorkqvist M, Wild EJ, Thiele J, Silvestroni A, Andre R, Lahiri N, et al. A novel pathogenic pathway of immune activation detectable before clinical onset in Huntington’s disease. J Exp Med. United States; 2008;205:1869–77.

15. Agus F, Crespo D, Myers RH, Labadorf A. The caudate nucleus undergoes dramatic and unique transcriptional changes in human prodromal Huntington’s disease brain. BMC Med Genomics. 2019;12:137.

16. Labadorf A, Hoss AG, Lagomarsino V, Latourelle JC, Hadzi TC, Bregu J, et al. Correction: RNA sequence analysis of human huntington disease brain reveals an extensive increase in inflammatory and developmental gene expression. PLoS One. 2016;11:1–21.

17. Durrenberger PF, Fernando FS, Kashefi SN, Bonnert TP, Seilhean D, Nait-Oumesmar B, et al. Common mechanisms in neurodegeneration and neuroinflammation: a BrainNet Europe gene expression microarray study. J Neural Transm. Austria; 2015;122:1055–68.

18. Langfelder P, Cantle JP, Chatzopoulou D, Wang N, Gao F, Al-Ramahi I, et al. Integrated genomics and proteomics define huntingtin CAG length-dependent networks in mice. Nat Neurosci. 2016;19:623–33.

19. Crotti A, Benner C, Kerman BE, Gosselin D, Lagier-Tourenne C, Zuccato C, et al. Mutant Huntingtin promotes autonomous microglia activation via myeloid lineage-determining factors. Nat Neurosci. 2014;17:513–21.

20. Audrain M, Haure-Mirande J-V, Wang M, Kim SH, Fanutza T, Chakrabarty P, et al. Integrative approach to sporadic Alzheimer’s disease: deficiency of TYROBP in a tauopathy mouse model reduces C1q and normalizes clinical phenotype while increasing spread and state of phosphorylation of tau. Mol Psychiatry. 2019;24:1383–97.

21. Haure-Mirande J-V, Wang M, Audrain M, Fanutza T, Kim SH, Heja S, et al. Integrative approach to sporadic Alzheimer’s disease: deficiency of TYROBP in cerebral Aβ amyloidosis mouse normalizes clinical phenotype and complement subnetwork molecular pathology without reducing Aβ burden. Mol Psychiatry. 2019;24:431–46.

22. Haure-Mirande J-V, Audrain M, Fanutza T, Kim SH, Klein WL, Glabe C, et al. Deficiency of TYROBP, an adapter protein for TREM2 and CR3 receptors, is neuroprotective in a mouse model of early Alzheimer’s pathology. Acta Neuropathol. 2017;134:769–88.

23. Zhang B, Gaiteri C, Bodea L-G, Wang Z, McElwee J, Podtelezhnikov AA, et al. Integrated systems approach identifies genetic nodes and networks in late-onset Alzheimer’s disease. Cell. 2013;153:707–20.

24. Turnbull IR, Colonna M. Activating and inhibitory functions of DAP12. Nat Rev Immunol. England; 2007;7:155–61.

25. Takaki R, Watson SR, Lanier LL. DAP12: an adapter protein with dual functionality. Immunol Rev. England; 2006;214:118–29.

26. Mócsai A, Abram CL, Jakus Z, Hu Y, Lanier LL, Lowell CA. Integrin signaling in neutrophils and macrophages uses adaptors containing immunoreceptor tyrosine-based activation motifs. Nat Immunol. 2006;7:1326–33.

27. Konishi H, Kiyama H. Microglial TREM2/DAP12 Signaling: A Double-Edged Sword in Neural Diseases. Front Cell Neurosci. 2018;12:206.

28. Bakker AB, Hoek RM, Cerwenka A, Blom B, Lucian L, McNeil T, et al. DAP12-deficient mice fail to develop autoimmunity due to impaired antigen priming. Immunity. United States; 2000;13:345–53.

29. Lier J, Streit WJ, Bechmann I. Beyond Activation: Characterizing Microglial Functional Phenotypes. Cells. 2021;10.

30. Fu R, Shen Q, Xu P, Luo JJ, Tang Y. Phagocytosis of microglia in the central nervous system diseases. Mol Neurobiol. 2014;49:1422–34.

31. Chistiakov DA, Killingsworth MC, Myasoedova VA, Orekhov AN, Bobryshev Y V. CD68/macrosialin: not just a histochemical marker. Lab Invest. United States; 2017;97:4–13.

32. Savage JC, St-Pierre M-K, Carrier M, El Hajj H, Novak SW, Sanchez MG, et al. Microglial physiological properties and interactions with synapses are altered at presymptomatic stages in a mouse model of Huntington’s disease pathology. J Neuroinflammation. 2020;17:98.

33. Hong S, Dissing-Olesen L, Stevens B. New insights on the role of microglia in synaptic pruning in health and disease. Curr Opin Neurobiol. 2016;36:128–34.

34. Wilton DK, Mastro K, Heller MD, Gergits FW, Willing CR, Frouin A, et al. Microglia Mediate Early Corticostriatal Synapse Loss and Cognitive Dysfunction in Huntington’s Disease Through Complement-Dependent Mechanisms. bioRxiv [Internet]. 2021;2021.12.03.471180. Available from: http://biorxiv.org/content/early/2021/12/04/2021.12.03.471180.abstract

35. Cepeda C, Hurst RS, Calvert CR, Hernández-Echeagaray E, Nguyen OK, Jocoy E, et al. Transient and progressive electrophysiological alterations in the corticostriatal pathway in a mouse model of Huntington’s disease. J Neurosci. 2003;23:961–9.

36. Gomez-Pastor R, Burchfiel ET, Neef DW, Jaeger AM, Cabiscol E, McKinstry SU, et al. Abnormal degradation of the neuronal stress-protective transcription factor HSF1 in Huntington’s disease. Nat Commun. 2017;8:14405.

37. Kim A, García-García E, Straccia M, Comella-Bolla A, Miguez A, Masana M, et al. Reduced Fractalkine Levels Lead to Striatal Synaptic Plasticity Deficits in Huntington’s Disease. Front Cell Neurosci. 2020;14:163.

38. Puigdellívol M, Cherubini M, Brito V, Giralt A, Suelves N, Ballesteros J, et al. A role for Kalirin-7 in corticostriatal synaptic dysfunction in Huntington’s disease. Hum Mol Genet. 2015;24:7265–85.

39. Langfelder P, Gao F, Wang N, Howland D, Kwak S, Vogt TF, et al. MicroRNA signatures of endogenous Huntingtin CAG repeat expansion in mice. PLoS One. 2018;13:1–20.

40. Novati A, Hentrich T, Wassouf Z, Weber JJ, Yu-Taeger L, Deglon N, et al. Environment-dependent striatal gene expression in the BACHD rat model for Huntington disease. Sci Rep. England; 2018;8:5803.

41. Brochier C, Gaillard M-C, Diguet E, Caudy N, Dossat C, Segurens B, et al. Quantitative gene expression profiling of mouse brain regions reveals differential transcripts conserved in human and affected in disease models. Physiol Genomics. United States; 2008;33:170–9.

42. Le Gras S, Keime C, Anthony A, Lotz C, De Longprez L, Brouillet E, et al. Altered enhancer transcription underlies Huntington’s disease striatal transcriptional signature. Sci Rep. England; 2017;7:42875.

43. Ament SA, Pearl JR, Grindeland A, St Claire J, Earls JC, Kovalenko M, et al. High resolution time-course mapping of early transcriptomic, molecular and cellular phenotypes in Huntington’s disease CAG knock-in mice across multiple genetic backgrounds. Hum Mol Genet. England; 2017;26:913–22.

44. Hervas-Corpion I, Guiretti D, Alcaraz-Iborra M, Olivares R, Campos-Caro A, Barco A, et al. Early alteration of epigenetic-related transcription in Huntington’s disease mouse models. Sci Rep. England; 2018;8:9925.

45. Becanovic K, Pouladi MA, Lim RS, Kuhn A, Pavlidis P, Luthi-Carter R, et al. Transcriptional changes in Huntington disease identified using genome-wide expression profiling and cross-platform analysis. Hum Mol Genet. England; 2010;19:1438–52.

46. Vuono R, Kouli A, Legault EM, Chagnon L, Allinson KS, La Spada A, et al. Association Between Toll-Like Receptor 4 (TLR4) and Triggering Receptor Expressed on Myeloid Cells 2 (TREM2) Genetic Variants and Clinical Progression of Huntington’s Disease. Mov Disord. 2020;35:401–8.

47. Collins BC, Hunter CL, Liu Y, Schilling B, Rosenberger G, Bader SL, et al. Multi-laboratory assessment of reproducibility, qualitative and quantitative performance of SWATH-mass spectrometry. Nat Commun. 2017;8:291.

48. Schilling B, Gibson BW, Hunter CL. Generation of High-Quality SWATH(®) Acquisition Data for Label-free Quantitative Proteomics Studies Using TripleTOF(®) Mass Spectrometers. Methods Mol Biol. 2017;1550:223–33.

49. Gillet LC, Navarro P, Tate S, Röst HSelevsek N, Reiter L, et al. Targeted data extraction of the MS/MS spectra generated by data-independent acquisition: a new concept for consistent and accurate proteome analysis. Mol Cell Proteomics. 2012;11:O111.016717.

50. Tennstaedt A, Pöpsel S, Truebestein L, Hauske P, Brockmann A, Schmidt N, et al. Human high temperature requirement serine protease A1 (HTRA1) degrades tau protein aggregates. J Biol Chem. 2012;287:20931–41.

51. Singhrao SK, Neal JW, Morgan BP, Gasque P. Increased complement biosynthesis by microglia and complement activation on neurons in Huntington’s disease. Exp Neurol. 1999;159:362–76.

52. Merienne N, Meunier C, Schneider A, Seguin J, Nair SS, Rocher AB, et al. Cell-Type-Specific Gene Expression Profiling in Adult Mouse Brain Reveals Normal and Disease-State Signatures. Cell Rep. 2019;26:2477-2493.e9.

53. Lee H, Fenster RJ, Pineda SS, Gibbs WS, Mohammadi S, Davila-Velderrain J, et al. Cell Type-Specific Transcriptomics Reveals that Mutant Huntingtin Leads to Mitochondrial RNA Release and Neuronal Innate Immune Activation. Neuron. 2020;107:891-908.e8.

54. Benraiss A, Mariani JN, Osipovitch M, Cornwell A, Windrem MS, Villanueva CB, et al. Cell-intrinsic glial pathology is conserved across human and murine models of Huntington’s disease. Cell Rep. United States; 2021;36:109308.

55. Seong IS, Woda JM, Song JJ, Lloret A, Abeyrathne PD, Woo CJ, et al. Huntingtin facilitates polycomb repressive complex 2. Hum Mol Genet. 2009;19:573–83.

56. Laugesen A, Højfeldt JW, Helin K. Molecular Mechanisms Directing PRC2 Recruitment and H3K27 Methylation. Mol Cell. 2019;74:8–18.

57. Ayata P, Badimon A, Strasburger HJ, Duff MK, Montgomery SE, Loh Y-HE, et al. Epigenetic regulation of brain region-specific microglia clearance activity. Nat Neurosci. 2018;21:1049–60.

58. Zhang H, Zhang T, Wang D, Jiang Y, Guo T, Zhang Y, et al. IFN-γ regulates the transformation of microglia into dendritic-like cells via the ERK/c-myc signaling pathway during cerebral ischemia/reperfusion in mice. Neurochem Int. England; 2020;141:104860.

59. Kaminska B, Mota M, Pizzi M. Signal transduction and epigenetic mechanisms in the control of microglia activation during neuroinflammation. Biochim Biophys Acta. Netherlands; 2016;1862:339–51.

60. Chen MJ, Ramesha S, Weinstock LD, Gao T, Ping L, Xiao H, et al. Extracellular signal-regulated kinase regulates microglial immune responses in Alzheimer’s disease. J Neurosci Res. United States; 2021;99:1704–21.

61. Bowles KR, Jones L. Kinase signalling in Huntington’s disease. J Huntingtons Dis. Netherlands; 2014;3:89–123.

62. Abd-Elrahman KS, Hamilton A, Hutchinson SR, Liu F, Russell RC, Ferguson SSG. mGluR5 antagonism increases autophagy and prevents disease progression in the zQ175 mouse model of Huntington’s disease. Sci Signal. United States; 2017;10.

63. Sanchis A, García-Gimeno MA, Cañada-Martínez AJ, Sequedo MD, Millán JM, Sanz P, et al. Metformin treatment reduces motor and neuropsychiatric phenotypes in the zQ175 mouse model of Huntington disease. Exp Mol Med. 2019;51:1–16.

64. Mukherjee S, Klaus C, Pricop-Jeckstadt M, Miller JA, Struebing FL. A Microglial Signature Directing Human Aging and Neurodegeneration-Related Gene Networks. Front Neurosci. 2019;13:2.

65. Mina E, van Roon-Mom W, Hettne K, van Zwet E, Goeman J, Neri C, et al. Common disease signatures from gene expression analysis in Huntington’s disease human blood and brain. Orphanet J Rare Dis. 2016;11:97.

66. Scarpa JR, Jiang P, Losic B, Readhead B, Gao VD, Dudley JT, et al. Systems Genetic Analyses Highlight a TGFβ-FOXO3 Dependent Striatal Astrocyte Network Conserved across Species and Associated with Stress, Sleep, and Huntington’s Disease. PLoS Genet. 2016;12:e1006137.

67. Neueder A, Bates GP. A common gene expression signature in Huntington’s disease patient brain regions. BMC Med Genomics. England; 2014;7:60.

68. Elorza A, Márquez Y, Cabrera JR, Sánchez-Trincado JL, Santos-Galindo M, Hernández IH, et al. Huntington’s disease-specific mis-splicing unveils key effector genes and altered splicing factors. Brain. 2021;144:2009–23.

69. Labadorf A, Hoss AG, Lagomarsino V, Latourelle JC, Hadzi TC, Bregu J, et al. RNA Sequence Analysis of Human Huntington Disease Brain Reveals an Extensive Increase in Inflammatory and Developmental Gene Expression. PLoS One. United States; 2015;10:e0143563.

70. Al-Dalahmah O, Sosunov AA, Shaik A, Ofori K, Liu Y, Vonsattel JP, et al. Single-nucleus RNA-seq identifies Huntington disease astrocyte states. Acta Neuropathol Commun. 2020;8:19.

71. Simmons DA, Casale M, Alcon B, Pham N, Narayan N, Lynch G. Ferritin accumulation in dystrophic microglia is an early event in the development of Huntington’s disease. Glia. United States; 2007;55:1074–84.

72. Paldino E, Balducci C, La Vitola P, Artioli L, D’Angelo V, Giampà C, et al. Neuroprotective Effects of Doxycycline in the R6/2 Mouse Model of Huntington’s Disease. Mol Neurobiol. 2020;57:1889–903.

73. Zeitler B, Froelich S, Marlen K, Shivak DA, Yu Q, Li D, et al. Allele-selective transcriptional repression of mutant HTT for the treatment of Huntington’s disease. Nat Med. United States; 2019;25:1131–42.

74. Diaz-Castro B, Gangwani MR, Yu X, Coppola G, Khakh BS. Astrocyte molecular signatures in Huntington’s disease. Sci Transl Med. United States; 2019;11.

75. Martín-Flores N, Pérez-Sisqués L, Creus-Muncunill J, Masana M, Ginés S, Alberch J, et al. Synaptic RTP801 contributes to motor-learning dysfunction in Huntington’s disease. Cell Death Dis. 2020;11:569.

76. Cepeda C, Hurst RS, Calvert CR, Hernández-Echeagaray E, Nguyen OK, Jocoy E, et al. Transient and progressive electrophysiological alterations in the corticostriatal pathway in a mouse model of Huntington’s disease. J Neurosci. 2003;23:961–9.

77. Kraft AD, Kaltenbach LS, Lo DC, Harry GJ. Activated microglia proliferate at neurites of mutant huntingtin-expressing neurons. Neurobiol Aging. United States; 2012;33:621.e17-33.

78. Liddelow SA, Guttenplan KA, Clarke LE, Bennett FC, Bohlen CJ, Schirmer L, et al. Neurotoxic reactive astrocytes are induced by activated microglia. Nature [Internet]. Nature Publishing Group; 2017;541:481–7. Available from: http://www.ncbi.nlm.nih.gov/pubmed/28099414%0Ahttp://www.pubmedcentral.nih.gov/articlerender.fcgi?artid=PMC5404890

79. Yang C-S, Yuk J-M, Shin D-M, Kang J, Lee SJ, Jo E-K. Secretory phospholipase A2 plays an essential role in microglial inflammatory responses to Mycobacterium tuberculosis. Glia. United States; 2009;57:1091–103.

80. Liu N, Zhuang Y, Zhou Z, Zhao J, Chen Q, Zheng J. NF-κB dependent up-regulation of TRPC6 by Aβ in BV-2 microglia cells increases COX-2 expression and contributes to hippocampus neuron damage. Neurosci Lett. Ireland; 2017;651:1–8.

81. Fei M, Wang H, Zhou M, Deng C, Zhang L, Han Y. Podoplanin influences the inflammatory phenotypes and mobility of microglia in traumatic brain injury. Biochem Biophys Res Commun. United States; 2020;523:361–7.

82. Kano S-I, Choi EY, Dohi E, Agarwal S, Chang DJ, Wilson AM, et al. Glutathione S-transferases promote proinflammatory astrocyte-microglia communication during brain inflammation. Sci Signal. 2019;12.

83. Lee M, Schwab C, McGeer PL. Astrocytes are GABAergic cells that modulate microglial activity. Glia. United States; 2011;59:152–65.

84. Kuhn SA, van Landeghem FKH, Zacharias R, Färber K, Rappert A, Pavlovic S, et al. Microglia express GABA(B) receptors to modulate interleukin release. Mol Cell Neurosci. United States; 2004;25:312–22.

85. Favuzzi E, Huang S, Saldi GA, Binan L, Ibrahim LA, Fernández-Otero M, et al. GABA-receptive microglia selectively sculpt developing inhibitory circuits. Cell. United States; 2021;184:4048-4063.e32.

86. Francelle L, Galvan L, Brouillet E. Possible involvement of self-defense mechanisms in the preferential vulnerability of the striatum in Huntington’s disease. Front Cell Neurosci [Internet]. 2014;8:1–13. Available from: http://journal.frontiersin.org/article/10.3389/fncel.2014.00295/abstract

87. Liu EA, Schultz ML, Mochida C, Chung C, Paulson HL, Lieberman AP. Fbxo2 mediates clearance of damaged lysosomes and modifies neurodegeneration in the Niemann-Pick C brain. JCI insight. 2020;5.

88. Inoue M, Yamada J, Aomatsu-Kikuchi E, Satoh K, Kondo H, Ishisaki A, et al. SCRG1 suppresses LPS-induced CCL22 production through ERK1/2 activation in mouse macrophage Raw264.7 cells. Mol Med Rep. 2017;15:4069–76.

89. Hodges A, Strand AD, Aragaki AK, Kuhn A, Sengstag T, Hughes G, et al. Regional and cellular gene expression changes in human Huntington’s disease brain. Hum Mol Genet. 2006;15:965–77.

90. Kuleshov M V, Jones MR, Rouillard AD, Fernandez NF, Duan Q, Wang Z, et al. Enrichr: a comprehensive gene set enrichment analysis web server 2016 update. Nucleic Acids Res. England; 2016;44:W90–7.

91. Young K, Morrison H. Quantifying Microglia Morphology from Photomicrographs of Immunohistochemistry Prepared Tissue Using ImageJ. J Vis Exp. 2018;

92. Livak KJ, Schmittgen TD. Analysis of relative gene expression data using real-time quantitative PCR and the 2(-Delta Delta C(T)) Method. Methods. United States; 2001;25:402–8.

93. Tarca AL, Bhatti G, Romero R. A comparison of gene set analysis methods in terms of sensitivity, prioritization and specificity. PLoS One. 2013;8:e79217.

94. Escher C, Reiter L, MacLean B, Ossola R, Herzog F, Chilton J, et al. Using iRT, a normalized retention time for more targeted measurement of peptides. Proteomics. 2012;12:1111–21.

95. Bruderer R, Bernhardt OM, Gandhi T, Xuan Y, Sondermann J, Schmidt M, et al. Optimization of Experimental Parameters in Data-Independent Mass Spectrometry Significantly Increases Depth and Reproducibility of Results. Mol Cell Proteomics. 2017;16:2296–309.

96. Burger T. Gentle Introduction to the Statistical Foundations of False Discovery Rate in Quantitative Proteomics. J Proteome Res. United States; 2018;17:12–22.

97. Rohart F, Gautier B, Singh A, Lê Cao K-A. mixOmics: An R package for ‘omics feature selection and multiple data integration. PLoS Comput Biol. 2017;13:e1005752.

98. Kamburov A, Wierling C, Lehrach H, Herwig R. ConsensusPathDB--a database for integrating human functional interaction networks. Nucleic Acids Res. 2009;37:D623–8.

99. Kamburov A, Pentchev K, Galicka H, Wierling C, Lehrach H, Herwig R. ConsensusPathDB: toward a more complete picture of cell biology. Nucleic Acids Res. 2011;39:D712–7.

100. Mattei D, Ivanov A, van Oostrum M, Pantelyushin S, Richetto J, Mueller F, et al. Enzymatic Dissociation Induces Transcriptional and Proteotype Bias in Brain Cell Populations. Int J Mol Sci. 2020;21.

101. Creus-Muncunill J, Badillos-Rodríguez R, Garcia-Forn M, Masana M, Garcia-Díaz Barriga G, Guisado-Corcoll A, et al. Increased translation as a novel pathogenic mechanism in Huntington’s disease. Brain. England; 2019;142:3158–75.

102. Cirnaru M-D, Creus-Muncunill J, Nelson S, Lewis TB, Watson J, Ellerby LM, et al. Striatal Cholinergic Dysregulation after Neonatal Decrease in X-Linked Dystonia Parkinsonism-Related TAF1 Isoforms. Mov Disord. United States; 2021;

