## Supplementary_materials for "Deletion of the microglial transmembrane immune signaling adaptor TYROBP ameliorates Huntington’s disease mouse phenotype": Supplementary_figures.pdf

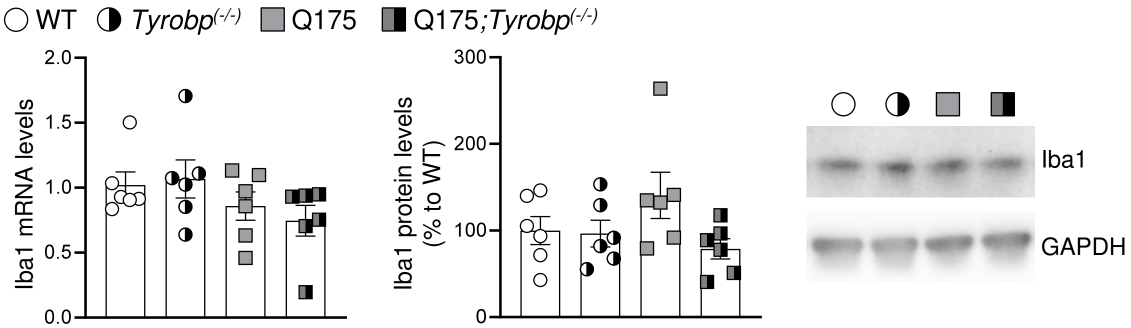


**Supplementary Fig. 1.** RT-qPCR of *Aif1* (Iba1 gene) mRNA (left) and WB of Iba1 protein (right) in the striatum of WT and Q175 mice with and without *Tyrobp* (10 months of age), n = 6 mice per group. Each point represents data from an individual mouse.

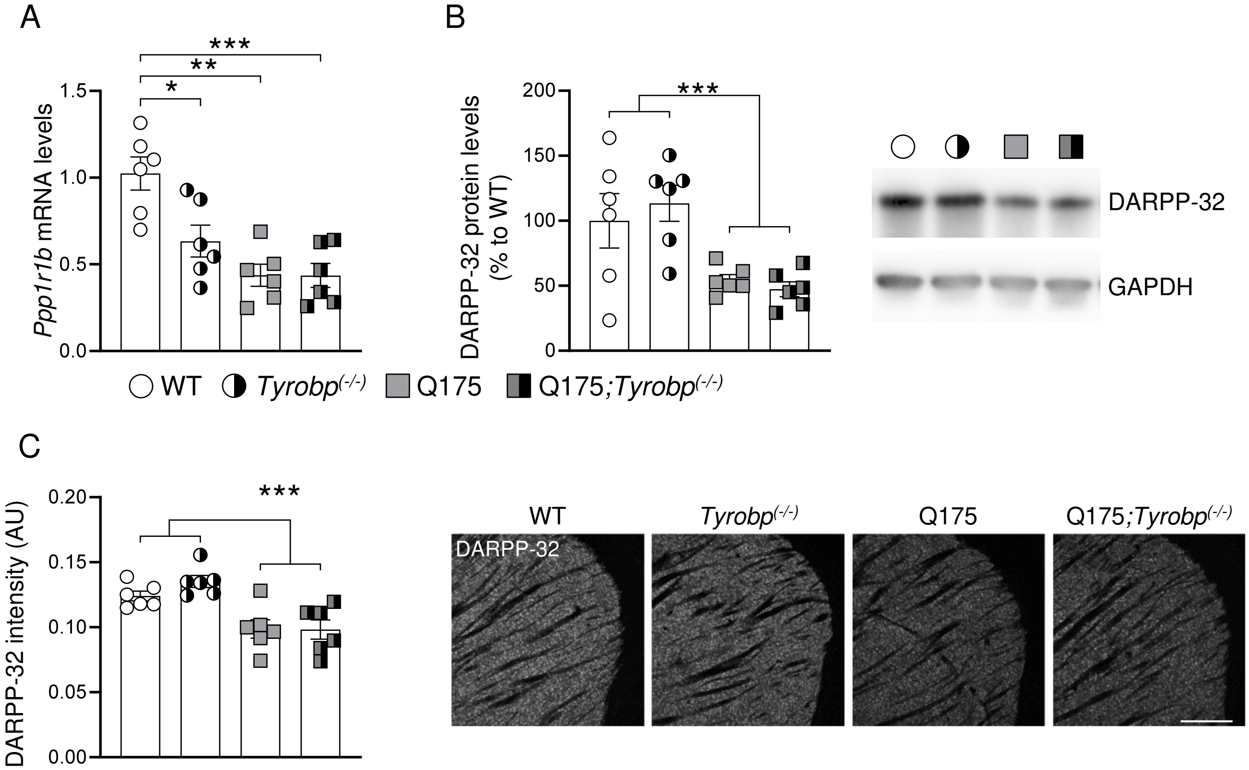


**Supplementary Fig. 2.** (A) RT-qPCR, (B) WB and (C) immunofluorescence analysis of DARPP-32 in the striatum of WT and Q175 mice with and without *Tyrobp* (10 months of age), n = 6 mice per group. Data represent the mean ± SEM. Each point represents data from an individual mouse. Two-way ANOVA followed by Bonferroni's post hoc test, *P < 0.05; **P < 0.01; ***P < 0.001. Scale bar = 200 µm.

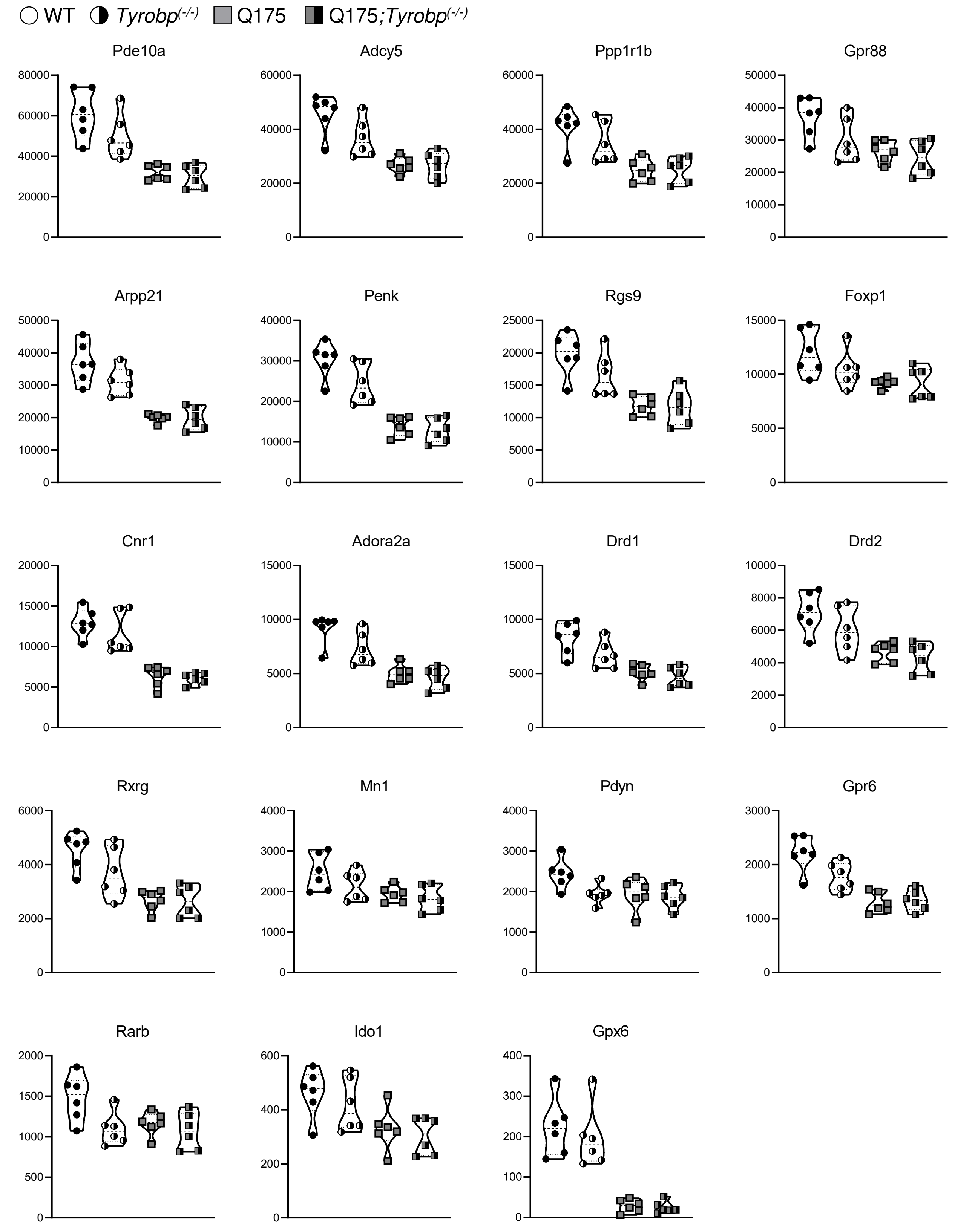


**Supplementary Fig. 3**. Normalized counts of striatal-specific genes detected in the striatum of WT and Q175 mice with and without Tyrobp (10 months of age) by bulk RNAseq, n = 6 mice per group.

A

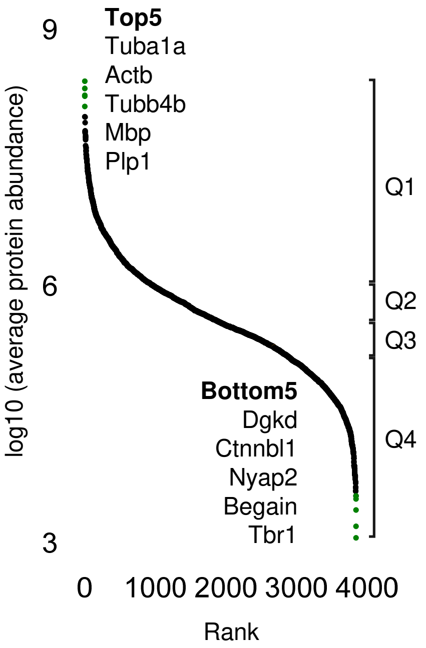


B

| Quartile | GO Biological Process 2021 | FDR | OR |
|---|---|---|---|
| Q1: | aerobic electron transport chain | 1.403E-35 | 33.1 |
| Q2: | cytoplasmic translation | 1.677E-22 | 14.26 |
| Q3: | post-Golgi vesicle-mediated transport | 4.992E-07 | 6.63 |
| Q4: | mitochondrial translational elongation | 2.689E-10 | 7.94 |

| Quartile | GO Cellular Component 2021 | FDR | OR |
|---|---|---|---|
| Q1: | focal adhesion | 2.217E-51 | 8.67 |
| Q2: | cytosolic large ribosomal subunit | 5.144E-14 | 14.61 |
| Q3: | late endosome | 3.342E-07 | 3.98 |
| Q4: | mitochondrial inner membrane | 0.0001579 | 2.67 |

**Supplementary Fig. 4**. (A) Ranking of brain proteins by normalized protein abundance from highest to lowest. (B) Top enrichment for each quartile is displayed for GO categories "biological process" and "cellular component"; FDR, Benjamini-Hochberg-corrected false discovery rate. OR, Odds Ratio.
